# Single Cell Analysis of Treatment–Resistant Prostate Cancer: Implications of Cell State Changes for Cell Surface Antigen Targeted Therapies

**DOI:** 10.1101/2024.04.09.588340

**Authors:** Samir Zaidi, Jooyoung Park, Joseph M. Chan, Martine P. Roudier, Jimmy L. Zhao, Anuradha Gopalan, Kristine M. Wadosky, Radhika A. Patel, Erolcan Sayar, Wouter R. Karthaus, D. Henry Kates, Ojasvi Chaudhary, Tianhao Xu, Ignas Masilionis, Linas Mazutis, Ronan Chaligné, Aleksandar Obradovic, Irina Linkov, Afsar Barlas, Achim Jungbluth, Natasha Rekhtman, Joachim Silber, Katia Manova–Todorova, Philip A. Watson, Lawrence D. True, Colm M. Morrissey, Howard I. Scher, Dana Rathkopf, Michael J. Morris, David W. Goodrich, Jungmin Choi, Peter S. Nelson, Michael C. Haffner, Charles L. Sawyers

## Abstract

Targeting cell surface molecules using radioligand and antibody–based therapies has yielded considerable success across cancers. However, it remains unclear how the expression of putative lineage markers, particularly cell surface molecules, varies in the process of lineage plasticity, wherein tumor cells alter their identity and acquire new oncogenic properties. A notable example of lineage plasticity is the transformation of prostate adenocarcinoma (PRAD) to neuroendocrine prostate cancer (NEPC)––a growing resistance mechanism that results in the loss of responsiveness to androgen blockade and portends dismal patient survival. To understand how lineage markers vary across the evolution of lineage plasticity in prostate cancer, we applied single cell analyses to 21 human prostate tumor biopsies and two genetically engineered mouse models, together with tissue microarray analysis (TMA) on 131 tumor samples. Not only did we observe a higher degree of phenotypic heterogeneity in castrate–resistant PRAD and NEPC than previously anticipated, but also found that the expression of molecules targeted therapeutically, namely *PSMA*, *STEAP1*, *STEAP2*, *TROP2, CEACAM5*, and *DLL3*, varied within a subset of gene–regulatory networks (GRNs). We also noted that NEPC and small cell lung cancer (SCLC) subtypes shared a set of GRNs, indicative of conserved biologic pathways that may be exploited therapeutically across tumor types. While this extreme level of transcriptional heterogeneity, particularly in cell surface marker expression, may mitigate the durability of clinical responses to novel antigen–directed therapies, its delineation may yield signatures for patient selection in clinical trials, potentially across distinct cancer types.

**SIGNIFICANCE STATEMENT:** Treatment of prostate cancer is rapidly evolving with several promising new drugs targeting different cell surface antigens. Selection of patients most likely to benefit from these therapies requires an understanding of how expression of these cell surface antigens varies across patients and how they change during disease progression, particularly in tumors that undergo lineage plasticity. Using immunohistochemistry and single cell mRNA sequencing, we reveal heterogeneity of cell states across a cohort of advanced disease prostate cancer patients; this heterogeneity is not captured by conventional histology–based designations of adenocarcinoma and neuroendocrine prostate cancer. We show these cell states can be identified by gene regulatory networks that could provide additional diagnostic precision based on their correlation with clinically relevant cell surface antigen expression.

## INTRODUCTION

In recent years, there has been a remarkable increase in the clinical development of antibody drug conjugates (ADCs), radioligand therapies (RLTs), bi–specific T cell engagers and chimeric antigen receptor expressing T cells (CAR–Ts), all of which are designed to target cell surface antigens expressed on cancer cells (*1–5*). ADCs selectively deliver potent chemotherapeutic toxins and RLTs deliver lethal doses of radiation, whereas bi–specifics and CAR–Ts leverage the immune system for tumor killing. All four approaches require expression of the target antigen on cancer cells (to ensure tumor reduction/elimination), and the level of expression often must be greater than in normal tissue to achieve an acceptable therapeutic index (*6, 7*). Consequently, clinical trials must be designed in a manner that ensures selection of patients that meet these criteria, often through a companion diagnostic. The development of a radiotheranostic for castrate–resistant prostate cancer (CRPC) is a particularly noteworthy example, wherein a small molecule for the prostate–specific membrane antigen (PSMA–617) is combined with a therapeutic radioisotope (^177^Lutetium) to specifically target prostate cancer cells (*1*).

Acquired resistance to conventional molecularly targeted therapies is often due to mutations within the drug target in rare clones that emerge during treatment (*8*). Many next generation inhibitors have been designed to overcome this form of “on target” resistance, with durable long–term remissions achieved in multiple tumor types including lung cancer (Osimertinib) and chronic myeloid leukemia (Asciminib) (*9*). However, a growing mode of resistance, commonly referred to as “lineage plasticity”, results from tumor cells adapting to environmental stresses, such as those associated with tumor invasion and metastases, as well as the selective pressure of drug therapy (*9–12*). The transition of adenocarcinoma to neuroendocrine cancer typifies this process and can be seen individually in up to 20% of prostate, lung, and gastric cancers who relapse on primary therapy. Unfortunately, cancer patients who harbor these plasticity–associated tumors have dismally short survival (*11, 13*).

To understand the repertoire of lineage states and, in that context, assess for cell surface marker expression across treatment–resistant prostate cancer, we utilized an integrated experimental and computational approach to analyze single cell RNA sequencing (scRNA-seq) from 21 human treatment–resistant prostate tumor biopsies and two genetically engineered mouse models (GEMMs), together with human tissue microarray analysis (TMA) comprising 131 CRPCs with prostate adenocarcinoma (PRAD) and neuroendocrine carcinoma (NEPC) histologies. The latter allowed spatial analysis at the protein level of lineage marker expression in tumors, including an assessment of inter– and intra–patient heterogeneity.

Through these comprehensive datasets, we find that PRAD and NEPC tumors display a high degree of phenotypic heterogeneity with an array of androgen receptor (AR) positive and negative, and NEPC gene regulatory networks (GRNs). Furthermore, through a comparative analysis of human small cell lung cancer (SCLC) and NEPC subtypes, we find a shared set of transcription factors (TFs) and cell surface antigens, indicative of conserved plasticity–associated gene programs. Lastly, by evaluating the expression of cell surface proteins that have been or are being targeted therapeutically, namely *PSMA, STEAP1/2, TROP2, CECAM5, and DLL3,* we find a high degree of heterogeneity within and across CRPC and NEPC patients and across different GRNs.

The degree of heterogeneity in the expression of cell surface markers in metastatic CRPC revealed by our analysis raises challenges in maximizing the clinical utility of cell surface targeted therapeutics in plasticity–associated states, underscoring the need to intervene prior to their emergence. Furthermore, the TF–specific signatures identified here could prove useful to more comprehensively classify patients, possibly across tumor types, based on evidence of shared regulatory networks.

## RESULTS

### Lineage Markers in Human Treatment–Resistant Prostate Cancer

To evaluate the fidelity of reported cell type and surface markers in treatment–resistant prostate cancer, we first performed immunohistochemistry on prostate cancer tissue microarrays (TMAs) (*14–16*) constructed from rapid autopsy samples of patients with advanced CRPC. The included cases span the clinical disease spectrum from adenocarcinoma to NEPC (**Methods**). The TMA consisted of 131 tumors from 16 patients, including primary prostate tissue and distant metastases, with 2 to 21 anatomically distinct tissue samples *per* patient. Samples were annotated by histology as PRAD, high–grade carcinoma (HGC) and NEPC (**Methods** for definitions), as well as by tumor site. Human TMAs were stained for the following markers: luminal or basal epithelial (AR, NKX3.1, CK8, and P63), neuroendocrine (SYP, INSM1, ASCL1, NEUROD1 and FOXA2), cell surface (TROP2 and DLL3), proliferation (KI67), as well as other markers of interest from scRNA–seq analyses of prostate cancer GEMMs (YAP1, POU2F3, MYC, SOX2, TFF3, and EZH2) (*17–19*) (**Figure 1A, Supplementary Table 1**). Levels of protein expression within histologies (*i.e* NEPC) were not significantly affected by ischemic time postmortem, except for *FOXA2* (**Supplementary Table 2**).

**Figure 1.**
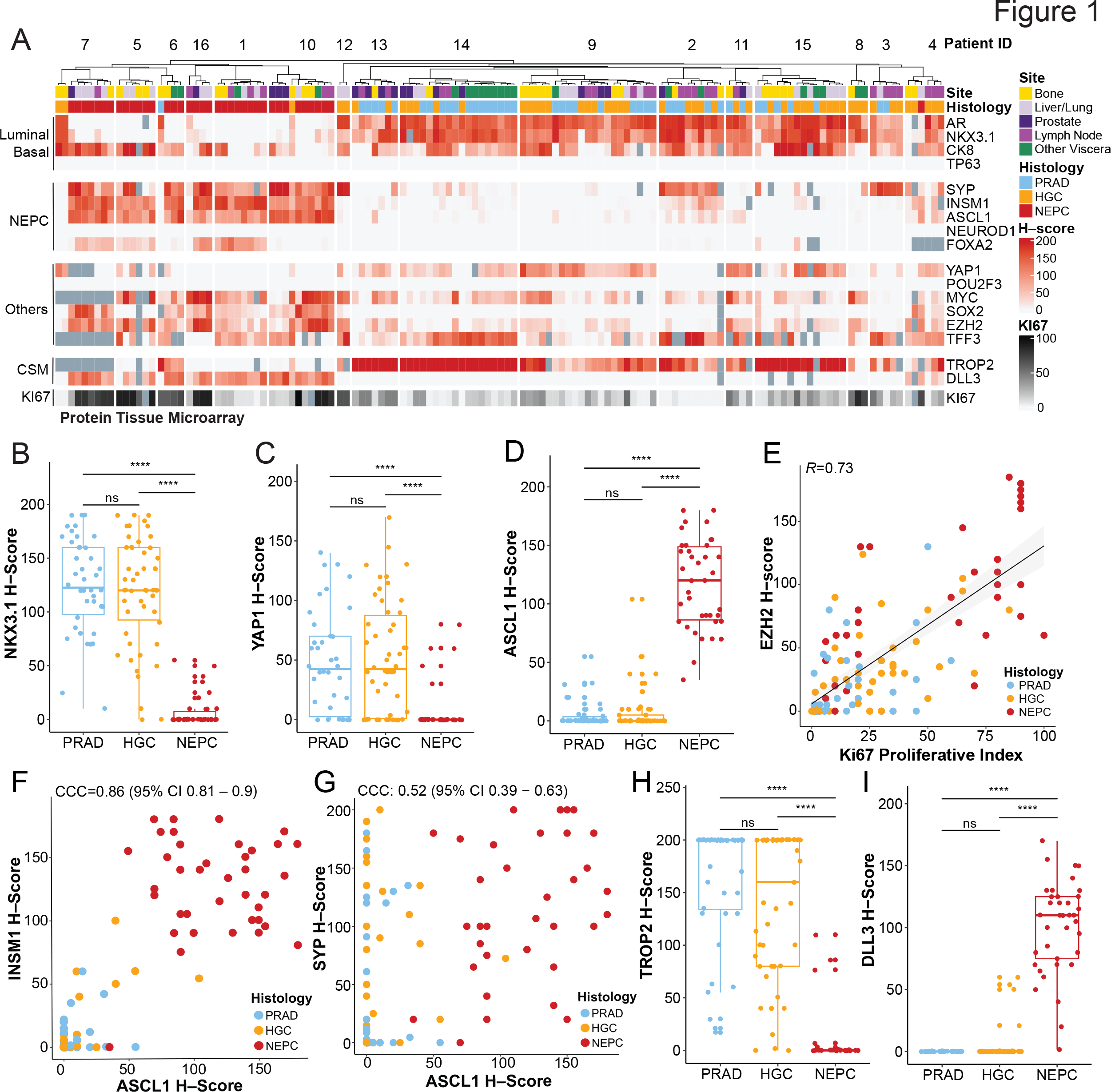
Tissue Microarray of Lineage and Cell Surface Markers in Human CRPC– adenocarcinoma and NEPC. (A) Heatmap of human CRPC tissue microarray–based immunohistochemical expression studies of patients from the rapid autopsy program at University of Washington. *H*-scores (immunohistochemical score, scale 0 to 200, and red gradient) are shown for select markers, namely luminal or basal (AR, NKX3.1, CK8, and P63), neuroendocrine prostate cancer (NEPC) (SYP, INSM1, ASCL1, NEUROD1, FOXA2), other single cell RNA– sequencing candidates from GEMMs (YAP1, POU2F3, CMYC, SOX2, EZH2, and TFF3), cell surface markers (CSM) (TROP2 and DLL3), and proliferative score (KI67, scale 0 to 100, and black gradient). Corresponding de–identified patient IDs (top row), site (bone, yellow; liver/lung, light purple; prostate, dark purple; lymph node, purple; other viscera, green), and histology (PRAD or prostate adenocarcinoma, light blue; HGC or high–grade carcinoma, orange; and NEPC or high–grade neuroendocrine, red) are labeled. Dark gray boxes are substituted in place of *H*– score for tumors with no immunohistochemical information. **(B–D)** Boxplot of H–scores of NKX3.1, YAP1, and ASCL1 grouped by histology (PRAD, HGC, and NEPC). Significance of H– score distribution was assessed by Wilcoxon signed–ranked test. **(E)** Scatter plot of H–scores of EZH2 (y–axis) and proliferative index of Ki67 (x–axis). Linear fit was calculated between two markers; the corresponding Pearson’s correlation is noted. **(F–G)** INSM1 or SYP (y–axis) and ASCL1 (x–axis) are shown with the color of the dot representing histology (PRAD, HGC, and NEPC) with corresponding Lin’s concordance correlation coefficient noted (95% confidence intervals). **(H–I)** Boxplot of H–scores of cell surface markers, TROP2 and DLL3 grouped by histology (PRAD, HGC, and NEPC). Of note, TROP2 and DLL3 expression has been assessed in a larger TMA (inclusive of these data) separated by categories: AR+/NE–, AR–/NE+, AR+/NE+, and AR–/NE– by our groups in Ajkunic *et al.* PMID 38296594. Significance of H–score distribution was assessed by Wilcoxon signed–ranked test. Abbreviations include: not significant (ns), * (<0.05), **(<0.01), ***(<0.001), ****(1x10^-4^).

As expected, PRADs showed high immunohistochemical scores for CK8, along with the prostate luminal markers AR (**Supplementary** Figure 1A) and NKX3.1 (**Figure 1B**) (*20*). This pattern was also noted in HGC but not in histologically classified NEPC tumors (example shown in **Supplementary** Figure 2A and 2D). The basal lineage marker P63 was absent in all tumors (**Figure 1A**, positive control shown in **Supplementary** Figure 1B). YAP1, a downstream nuclear effector of the Hippo pathway, has been implicated in the stem–cell like subsets of human tumoroids (*21*). YAP1 and ASCL1 H–scores showed an orthogonal relationship. Specifically, YAP1 was high in PRAD/HGC but not NEPC (**Figure 1C**, *P*<1x10^-4^), while ASCL1 and other neuroendocrine–associated TFs were high in NEPC histologies but not PRAD/HGC (**Figure 1D** and **Supplementary** Figure 1C). This profile mirrors that of YAP1 expression in lung adenocarcinoma *versus* small cell lung cancer (SCLC) (*22, 23*). Lastly, EZH2, a subunit of the polycomb repressive complex 2, which has been implicated in NEPC transformation (*24, 25*), showed higher expression in HGC and NEPC compared with PRAD (highest in NEPC) (**Supplementary** Figure 1A) and showed a positive correlation with the Ki–67 index (*R*=0.73, **Figure 1E**).

We next focused on additional neuroendocrine markers, beyond ASCL1, that have been previously described in SCLC and NEPC (*17, 23, 26*). Histologically defined NEPC tumors were enriched for expression of INSM1, SYP, FOXA2, and SOX2 (**Supplementary** Figure 1A, example of NEPC IHC stains shown in **Supplementary** Figure 2A and 2B). The H–score for INSM1 showed strong concordance with ASCL1 (CCC=0.86, 95% CI 0.81–0.9, **Figure 1F**) whereas SYP, often considered a canonical NEPC marker (*27*), was less concordant with ASCL1 (CCC=0.52, 95% CI 0.39–0.63) (**Figure 1G**) and other NEPC TFs (CCC=0.46, 95% CI 0.34–0.56) (**Supplementary** Figure 1D). This is likely because several SYP–positive tumors were negative for ASCL1 and INSM1 but positive for luminal markers such as AR, NKX3.1 and CK8. These AR–positive, SYP–positive tumors (often referred to as amphicrine, example shown **Supplementary** Figure 2C) have adenocarcinoma histology and clinically behave differently than bona fide NEPC tumors (*28*). These examples of SYP-positive, ASCL1–negative tumors suggest that SYP expression alone may not be sufficient to diagnose NEPC (*28, 29*).

Nuclear staining of NEUROD1, which marks a distinct small cell subtype in SCLC (*30*), was not detected in this human TMA cohort (**Figure 1A**, positive control stain shown in **Supplementary** Figure 1B). However, a NEUROD1–expressing NEPC subset has been implicated previously in ATAC–seq analysis of prostate cancer PDX models (*31*) and was detected by scRNA–seq in at least one NEPC sample not represented on the TMA (discussed below). This may be due to the overall low incidence of NEUROD1 expression in NEPC. POU2F3 expression was also neither detectable in the TMA by IHC (positive control stain shown in **Supplementary** Figure 1B), nor in the scRNA-seq cohort discussed below. Although clearly detectable in prostate GEMMs with NEPC histology, POU2F3 expression may be rare in human prostate cancers (*17, 18*). In contrast, TFF3, a mucosal–associated protein that is expressed in a subset of SCLCs and marks a non–neuroendocrine prostate population in prostate GEMMs (*10, 32*), was readily detected in human TMA specimens within subsets of PRAD, HGC and NEPC samples (**Supplementary** Figure 1A).

We next focused on the cell surface antigens TROP2 and DLL3, which are targets of various therapeutic agents currently in clinical trials. TROP2, the target of an FDA–approved antibody–drug conjugate in breast and bladder cancers (*33, 34*), was expressed in all PRADs and HGCs but not in the majority of NEPC samples (**Figure 1H**) nor in cells with expression of NEPC TFs (**Supplementary** Figure 1E). Conversely, expression of DLL3, the target of multiple agents currently under clinical investigation in SCLC (antibody drug conjugates, T cell engagers, CAR– T cells) (*35, 36*), was restricted to NEPC tumors (**Figure 1I**) and showed strong concordance with NEPC TFs (CCC=0.9, 95% CI 0.87 – 0.93) (**Supplementary** Figure 1F). This is consistent with a prior study from our groups documenting the expression of TROP2 and DLL3 in rapid autopsy samples (*37*).

Finally, the availability within the TMA to interrogate multiple independent metastatic sites from the same patient allowed us to detect intra–patient lineage heterogeneity. The most striking example from this analysis was expression of luminal epithelial markers (AR, NKX3.1, CK8) within individual bone or soft tissues metastases of three patients (Patients 6,10, and 16) with a clinical diagnosis of NEPC based on analysis of other tissue sites. These site-specific lineage differences are consistent with the notion that tissue microenvironmental signals may influence lineage conversion (*10*) (**Figure 1A** and **Supplementary Table 1** for patient–specific TMA H–scores).

Taken together, profiling of late stage CRPC with a broad panel of lineage markers documents that (a) YAP1 loss generally occurs in ASCL1–positive NEPC tumors, (b) TROP2 is predominantly expressed in PRAD/HGC, whereas DLL3 is almost exclusively present in NEPC tumors, (c) in comparison to ASCL1, the expression of other transcription factors linked with neuroendocrine phenotypes, such as NEUROD1 and POU2F3, is less common, and (d) SYP expression alone has limitations as a diagnostic marker for NEPC. Furthermore, our TMA analysis demonstrates that a single–site biopsy is insufficient to adequately capture the intra– tumoral heterogeneity in late–stage prostate cancer patients.

### Diverse Transcriptional Networks in Human CRPC

To extend our analysis of lineage heterogeneity in human CRPC beyond *in situ* methods, we studied gene expression networks in a set of human tumor biopsies through single cell RNA sequencing. We previously reported the transcriptomic architecture of 12 CRPC biopsies to identify JAK–STAT and FGFR as signaling pathways required for plasticity (*10*). We now report gene regulatory networks (GRNs) on an expanded cohort of 23 tumors (from 21 unique patients), including 9 naïve or castration–sensitive prostate cancer (*38*) and 14 late–stage metastatic CRPC tumors (119,083 profiled cells) (*10*). All tumors were reviewed by a genitourinary pathologist and classified histologically as CRPC–adenocarcinoma or NEPC. Furthermore, all CRPC– adenocarcinoma or NEPC tumors had been treated with more than two lines of therapy at the time of biopsy, with the majority having received taxanes (refer to **Supplementary Table 3** for details on histology, tissue site, tumor genomics and prior treatment, and **Figure 2A**).

**Figure 2.**
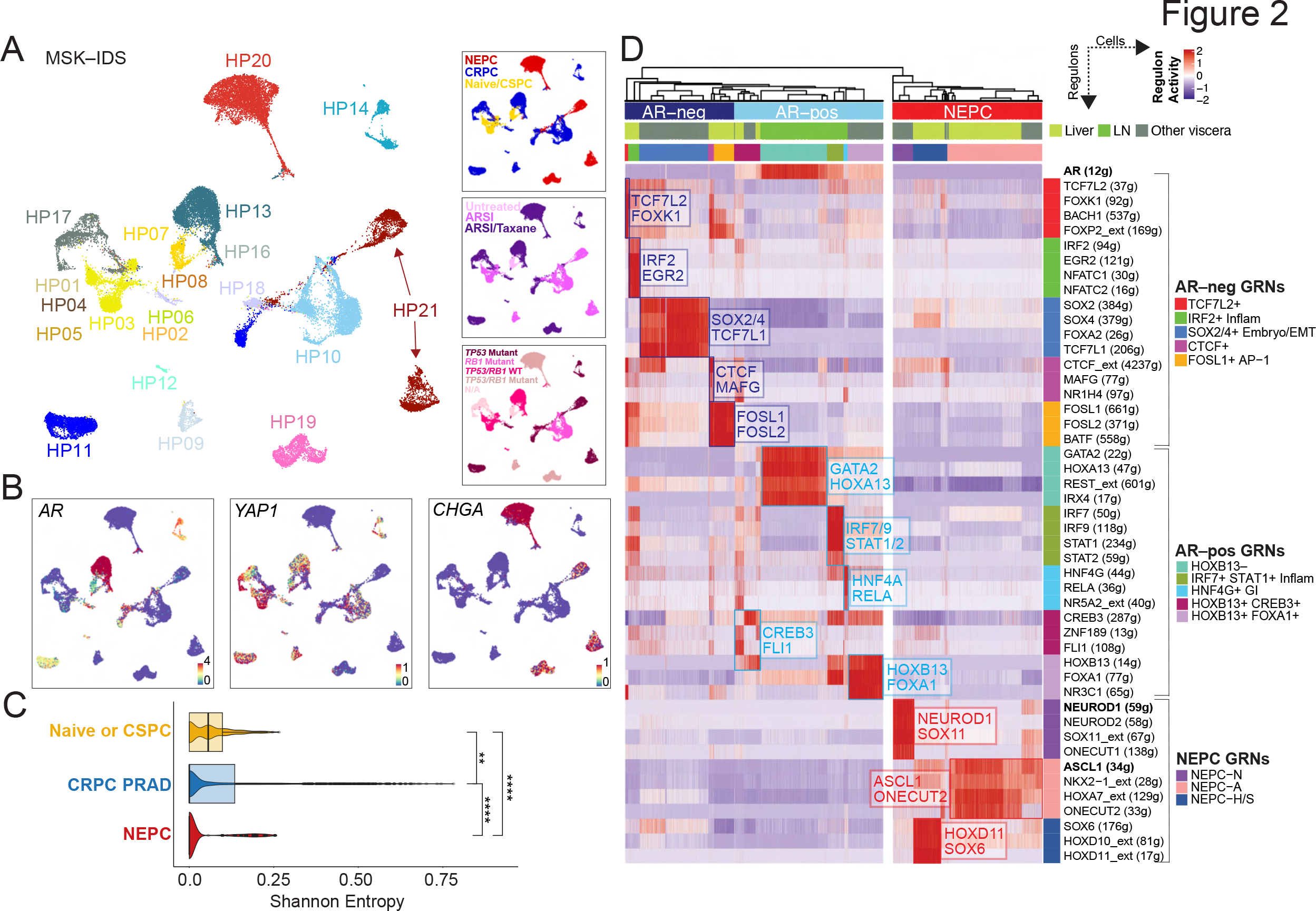
Diverse Gene–Regulatory Networks in Castration–Resistant Prostate Cancer. (A) UMAP of tumor cells (N=35,696 cells), colored by patient ID (large panel on left), category (top right panel), treatments (middle right panel; categories include untreated, androgen–receptor signaling inhibitor/ARSI, and ARSI plus taxane–based chemotherapy) or *TP53/RB1* genomic status (bottom right panel). Also detailed in Supplementary Table 3. **(B)** UMAPs showing expression [log(X +1)] of lineage genes, namely *AR*, *YAP1*, and *CHGA*. **(C)** Boxplot of inter– patient heterogeneity measured by Shannon entropy based of patient frequencies. To control for cell sampling, 100 cells were subsampled from each Phenograph cluster (*k*=30) within tumor compartments 100 times with replacement (Wilcoxon signed–rank test, Methods). Immune and mesenchymal inclusion shown in Supplementary Figure 3H. Abbreviations: * (<0.05), ****(1x10^-4^). **(D)** Heatmap of CRPC–adenocarcinoma and NEPC cells (x–axis) and *per* cell scaled regulon activity scores (z–score: –2 to 2) is shown for select TFs (paratheses denotes number of genes within regulon, extended heatmap in Supplementary Figure 4). A dendrogram cutoff of 15 based on adjusted Rand index was used to unbiasedly define the number of gene–regulatory networks (GRNs), yielding 10 and 3 CRPC–adeno and NEPC GRNs, respectively. Regulons were assigned to GRNs based on regulon specificity score (RSS) and ranked by significance (Supplementary Table 6). Adenocarcinoma GRNs were labeled based on AR activity (light blue on top panel of heatmap; bracketed by AR–positive GRNs) and without or having low AR activity (dark blue on top panel of heatmap; bracketed by AR–negative GRNs). NEPC regulons are shown (red on top panel of heatmap; bracketed by NEPC GRNs). *AR(12g)*, *NEUROD1(59g)*, and *ASCL1(34g)* regulons are **bolded** for reference.

Unsupervised clustering was used to iteratively label coarse cell types into lineage– defined groups using canonical markers (**Methods**) (**Supplementary** Figure 3A–E **and Supplementary Table 4**). 35,696 primary naïve or CSPC and metastatic CRPC tumor cells were labeled using select tumor markers (**Figure 2B** and **Supplementary** Figure 3B and C) and copy number detection (**Supplementary** Figure 3D). Given the observation of lineage marker heterogeneity in the CRPC TMA, we assessed the degree of inter–patient heterogeneity by calculating the Shannon diversity of different patient phenotypes (**Methods**). Clusters of cells associated with CRPC PRAD and NEPC were significantly more heterogenous (patient–specific, lower entropy) than those from CSPC tumors (**Figure 2C** and **Supplementary** Figure 3F), consistent with the notion that plasticity arises after androgen deprivation therapy (*11*). Furthermore, compared with tumor cells, higher entropy (lower phenotypic diversity or multi– patient phenotypes) was noted in stromal, myeloid and lymphoid cell populations (**Supplementary** Figure 3G–H), as has been described in single cell analyses of other cancer types (*32, 39*).

To study tumor cell heterogeneity specifically in CRPC PRAD and NEPC samples and reasoning that lineage plasticity is likely driven by transcription factor (TF) networks, we focused on shared and unique gene–regulatory networks (GRNs) across samples using single–cell regulatory network inference (SCENIC), (**Figure 2D, Supplementary** Figure 4A–C, and **Supplementary Table 5**). SCENIC has been utilized effectively to identify GRNs and cell types from single cell RNA–sequencing data with improved accuracy when integrated with chromatin accessibility data (*40, 41*). We thus used hierarchical clustering of regulon activity within tumor cells, which unbiasedly identified 10 distinct putative GRNs in CRPC PRAD and 3 GRNs in NEPC tumor cells (**Methods,** refer to cell– and patient–based robustness analyses for recurrent GRNs in **Supplementary** Figure 5A**/5B and 6**, respectively). We further ranked regulons for differential activity within each GRN (**Supplementary Table 6**) (**Methods**). The 10 CRPC adenocarcinoma GRNs broadly separated based on activity of the *AR* regulon, the dominant oncogenic pathway in CRPC. There were five GRN groups with *AR* regulon activity. Two of the identified *AR*+ GRNs displayed high *HOXB13* activity, showing either higher levels of *FOXA1* (labeled as *AR*+*HOXB13*+*FOXA1*+) or CREB3 (labeled as *AR*+*HOXB13*+*CREB3*+). One *AR*+ regulon showed lower *HOXB13* and higher *GATA2* and *HOXA13* activity (labeled as *AR*+*HOXB13*–), largely derived from one sample (MSK–HP13) that had lost *FOLH1* (which encodes for PSMA) expression. The lower expression of *HOXB13* is consistent with recent reports implicating *HOXB13* as a potential regulator of *PSMA* expression (*42*). We identified two further *AR*+ GRNs: “inflammatory” displaying high activities for *IRF7*/*9* and *STAT1*/*2* (*10*) (labeled as *AR*+ *IRF7*+*STAT1*+ Inflam) and “GI–lineage” with high activity of *HNF4G* and *RELA (AR+HNF4G*+ GI). The GI–lineage regulon showed enrichment of a mid–to–hindgut differentiation pathway, consistent with prior studies describing a role for *HNF4G* in promoting castration resistance (**Supplementary Table 7**) (*43*).

The other five CRPC PRAD GRNs identified by hierarchical clustering lacked or had low activity of the *AR* regulon. One of these included activity for *SOX2* and *SOX4*, along with *FOXA2*, *TCF7L1*, and *TWIST2* (**Figures 2D** and **Supplementary Table 6**) (labeled as *SOX2/4*+ Embryo/EMT). These genes are highly expressed in the developing embryo and are enriched in the epithelial–to–mesenchymal transition and WNT signaling (*TCF4*) (**Supplementary Table 7**). Another GRN had high activity for *IRF2*, along with *NFATC1*/*2* and *EGR2*, consistent with our recent report of tumor–intrinsic inflammatory JAK/STAT signaling and inflammatory programs driving lineage plasticity (labeled as *IRF2*+ Inflam) (*10*). *BATF* and *FOSL1*/*2* marked another non–AR GRN, concordant with a stem–cell–like group identified from patient–derived tumoroids and xenografts (FOSL1/2+ AP–1) (*21*) (**Figures 2D and Supplementary Table 6**). These latter cells were enriched for stem cell programs and the AP–1 pathway based on GSEA (**Supplementary Table 7**). Two GRNs, *albeit* comprising a smaller number of tumor cells, showed high activity for the *TCF7L2* regulon (along with *KLF8*, *FOXK1*, *FOXP2*, *BACH1*) (labeled TCF7L2+) and *CTCF* (along with MAFG and NR1H) (labeled CTCF+) respectively (**Figures 2D** and **Supplementary Table 6**). *TCF7L2* was previously identified as the top transcription factor (TF) candidate that marks a WNT–dominant CRPC phenotype (*21*).

Finally, we noted three putative NEPC GRNs largely distinguished by *ASCL1* and *NEUROD1* (*31*). NEPC–*ASCL1*+ cells (NEPC–A) showed high expression of *E2F* and neuronal targets (**Supplementary Table 7**), along with *ONECUT2* and *NKX2–1* (**Figure 2D**). Within a population of cells with lower ASCL1 activity, there was a subgroup that showed higher activity of *HOXD11* and *SOX6* (NEPC–H/S) (**Supplementary Table 6**) that was also enriched for NOTCH and β–catenin signaling (**Supplementary Table 7**). Within the *NEUROD1* GRN (NEPC–N), there was activity for *NEUROD2*, *ONECUT1*, *and SOX11* (**Figure 2D**). This group is akin to SCLC–N and showed strong overrepresentation of BMP signaling (*SOX11* and *ZNF423*) (**Supplementary Table 7**).

### Convergence Between CRPC, SCLC and GEMM Regulons

Given that the aforementioned analysis of human CRPC tumors yielded snapshots into an array of CRPC PRAD and NEPC transcriptional states, we re–analyzed previously published single cell sequencing data from GEMMs across multiple time points during the adenocarcinoma– to–neuroendocrine transition (*10*). This allowed us not only to identify overlapping TFs between mouse and human tumors but, importantly, to also detect potential intermediate cell populations that may not be captured in snapshots of human tumors. In our GEMM models of prostate– specific deletion of *Pten*, *Rb1* and/or *Tp53*, we focused on two genotypes at varying time points of tumorigenesis. PtR mice (*Pb*-*Cre*;*Pten^flox/flox^*;*Rb1^flox/flox^*) were studied at 24, 30 and 47 weeks, and PtRP mice (*Pb*-*Cre*;*Pten^flox/flox^*;*Rb1^flox/flox^*;*Tp53^flox/flox^*) at 8, 9, 12 and 16 weeks (*10*).

By implementing SCENIC, we found nine tumor–associated GRNs within the PtR and PtRP GEMMs including one defined by *Stat1/2* and *Irf2/7/9*, validating our recent findings on the critical role of JAK/STAT signaling in initiating plasticity (**Figure 3A** and **3B, Supplementary** Figure 7, and **Supplementary Table 8–9**). Furthermore, certain human CRPC and putative GEMM GRNs showed an overlap of specific cell populations, including NEPC–A and the above mentioned inflammatory GRN with high activity of *IRF7*/*9* and *STAT1*/*2* (**Supplementary** Figure 8A**/B**).

**Figure 3.**
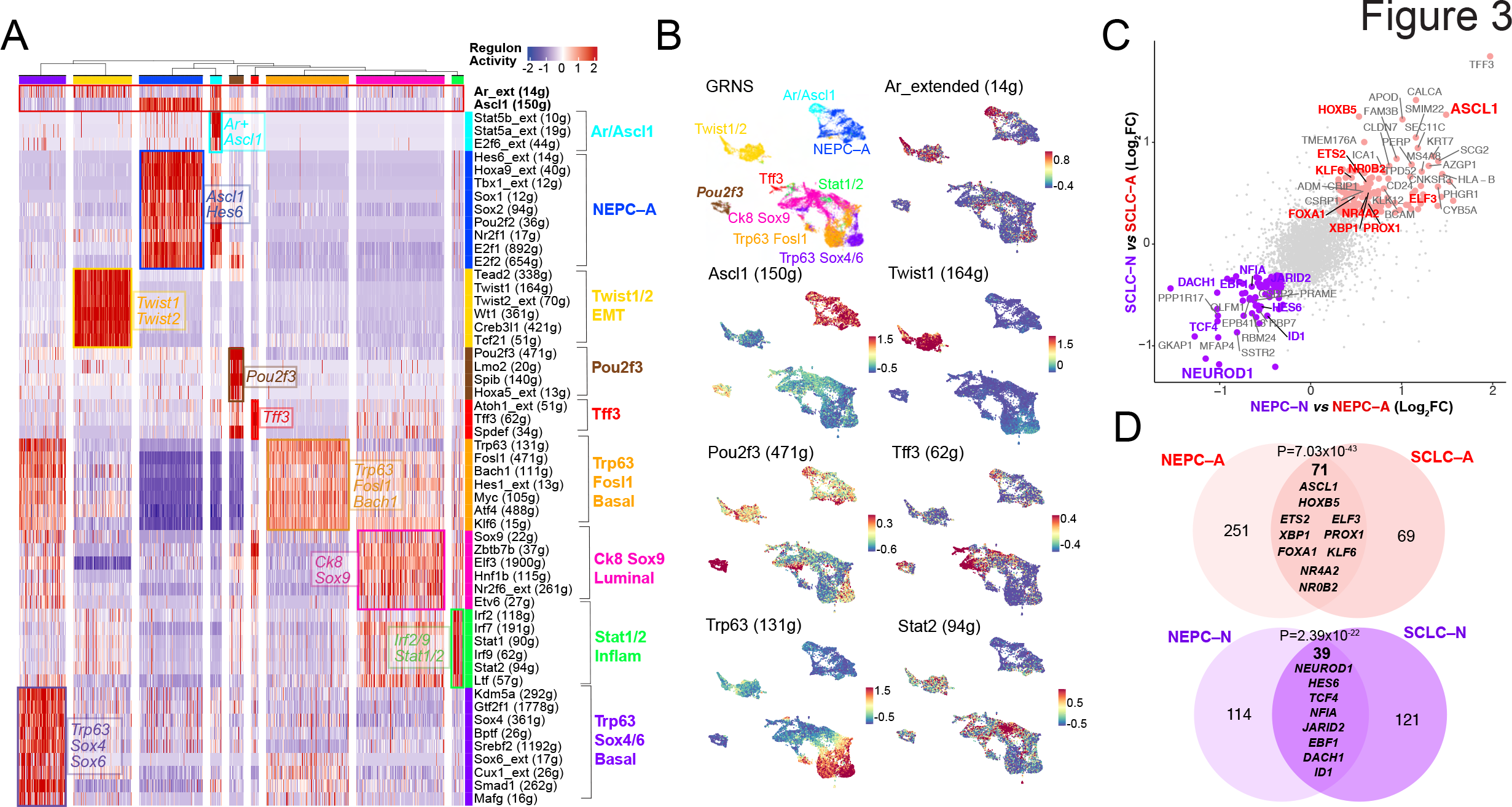
GEMM GRNs and NEPC and SCLC Overlap. (**A**) Heatmap of GEMM tumor cells (*N*=21,499) (x–axis) and and *per* cell scaled regulon activity scores (z–score: –2 to 2) is shown for select TFs (paratheses denotes number of genes within regulon). A dendrogram cutoff of 12 based on adjusted Rand index yielded 9 GRNs with regulons assigned to GRNs based on regulon specificity score (RSS) and ranked by significance (Methods, Supplementary Table 9). *Ar*– extended (14g) and *Ascl1* (150g) are shown in the top, bolded, and boxed in red for reference. (**B**) UMAP of GEMMs mutant *Gfp*–positive cells are colored by annotated GRN (color scheme corresponds to in Figure 3A), or by regulon activity (z–score) of *Ar_extended (14g), Ascl1 (150g), Twist1 (164g), Pou2f3 (471g), Tff3 (62g), Trp63 (131g), and Stat2 (94g).* (**C**) NEPC–N *vs.* NEPC– A (shown on x–axis) or SCLC–N *vs.* SCLC–A (shown on y–axis) were compared using MAST and the log2FC for each gene is shown on the scatter plot. Genes with log2FC > 0.4 (and padj<0.05) are labeled with TFs noted in red or purple for being enriched in both NEPC and SCLC *ASCL1* and *NEUROD1* subsets, respectively. (**D**) Venn diagram shows the overlap of top DEGs (average log2FC > 0.4, adjusted p–value < 0.05) shared between NEPC–A and SCLC–A (red) or NEPC–N and SCLC–N (purple). A Fisher’s exact test was used for significance of overlap.

This comparative analysis also allowed the detection of GRNs unique to GEMMs that may represent intermediate states. Specifically, *2* GRNs showed activity of the basal cell lineage factor *Trp63*, together with the co–expression of *Hes1, Bach1, and Fosl1*. One of the *Trp63*–marked clusters also displayed higher levels of *Ar* activity and high regulon activity for *Sox4, Sox6* and *Cux1*; the latter TFs have been implicated in dendritogenesis and neuronal differentiation (*44*) (**Figure 3A** and **3B**, and **Supplementary Table 10**). Given the lack of P63 in the human TMA and human scRNA–sequencing dataset, P63–positive tumors likely represent a rare entity, or an intermediary state not readily captured in human tumors. This is in comparison to our detection of P63-negative basal–like populations in human tumors, consistent with prior reports and as shown in our single cell dataset (**Supplementary** Figure 3E). In addition, we previously reported a unique non–neuroendocrine population marked by the tuft cell marker *Pou2f3* (**Figure 3A** and **3B**); however, we have been unable to find convincing evidence of POU2F3 expression in human CRPC at the RNA or protein level, as discussed earlier (*17*). Lastly, one smaller subset with Ascl1 expression also displayed *Ar* and *Stat5a/5b* activity (Ar/Ascl1), suggesting that *Ar* and its regulon may be expressed within a subset of *Ascl1*–positive NEPC GEMM cells (**Figure 3A** and **3B**). Note this GRN is distinct from the AR–positive, SYP–positive (amphicrine) but ASCL1–negative human CRPC tumors discussed earlier (**Figure 1A**).

Given the consistent presence of *ASCL1* GRNs in both human CRPC and GEMM NEPCs, we utilized SCLC data to determine whether there are common transcriptional networks across prostate and lung histologies (*32*). We found significant enrichment of NEPC–A and NEPC–N regulons in corresponding SCLC–A and SCLC–N subsets, respectively (**Figure 3C** and **3D, Supplementary** Figure 9A, and **Supplementary Table 11**). Examples of such shared TFs between NEPC and SCLC subtypes included: *ASCL1, HOXB5*, *ETS2*, *ELF3*, *XBP1 and PROX1* (*ASCL1* subtype), and *NEUROD1*, *HES6*, *TCF4, NFIA,* and *JARID2* (*NEUROD1* subtype).

Furthermore, while we did not detect the GEMM *Pou2f3* subset in our human dataset, we compared this GRN with SCLC–P, and noted that both the GEMM *Pou2f3* and inflammatory GRNs showed enrichment in SCLC–P. There were TFs, namely *POU2F3*, *SMARCC1,* and *MYB*, which were enriched in both *Pou2f3* and SCLC–P populations (**Supplementary** Figure 9B–D).

### Targeting Lineage Plasticity States

The recent approval of PSMA–targeted therapies has directed attention into the degree of inter– and intra–patient heterogeneity, particularly as it may impact therapeutic response (*15*). Given our identification of putative GRNs in both murine and human treatment–resistant tumors, we explored the expression of several prostate cancer targets in our dataset, notably *FOLH1 (PSMA), STEAP1/STEAP2, TACSTD2 (TROP2*)*, CEACAM5 and DLL3*, all of which have drug candidates in various stages of clinical development (*1–5*).

We first focused on *PSMA* given the expanding clinical usage of Lu^177^–PSMA–617 (Pluvicto) for advanced CRPC (*1*). Upon scoring each regulon for its average expression of *FOLH1/PSMA*, we noted a positive association with most AR–positive GRN within the CRPC PRAD samples (CCC=0.71) (**Figure 4A**). The highest ranking *PSMA*^high^/*AR*^high^ GRNs were associated with the luminal *HOXB13+* or inflammatory *IRF7/9+* GRNs (**Figures 4A** and **4B**). In contrast, the *AR+* GI GRN with *HNF4G* activity had lower expression of both *PSMA* and *AR*. There was also a *PSMA^l^*^ow^/*AR*^high^ GRN, from patient MSK–HP13, which had lower *HOXB13, KLF15, NFIL3* activity, but showed the highest activity for *AR*, and was enriched for *GATA2*, *HOXA6*, *HOXA13*, and *RELB* (**Figure 4B** and **Supplementary** Figure 10A–B). There were several *PSMA*^low^/*AR*^low^ CRPC GRNs that were enriched for genes and pathways related to embryonic, epithelial–to–mesenchymal and/or WNT pathways (**Supplementary Table 12**). Furthermore, NEPC samples within our cohort did not express PSMA and, as expected, displayed minimal AR signaling activity (**Figure 4A**); however, it is possible that aberrant PSMA expression may be present in NEPC given reports of its co–expression with *HOXB13* (*42*). Lastly, we analyzed each tumor sample for intra–tumoral PSMA heterogeneity. Patient MSK–HP13 showed a cluster of *AR*–positive, *PSMA*–positive cells, while the remaining clusters were *AR*–positive, *PSMA*–negative (**Figure 4C**). Certain TFs followed this pattern of negative PSMA expression with AR–positivity, including *HOXB13*, *SOX4*, and *GATA2* **(Supplementary** Figure 10C). Whether the high level of PSMA expression in a subset of cells from this lesion is sufficient to score as PSMA–positive on a PET scan is unknown, but cases such as this underscore the potential of heterogeneous PSMA expression even within single tumor foci (*15, 45*).

**Figure 4.**
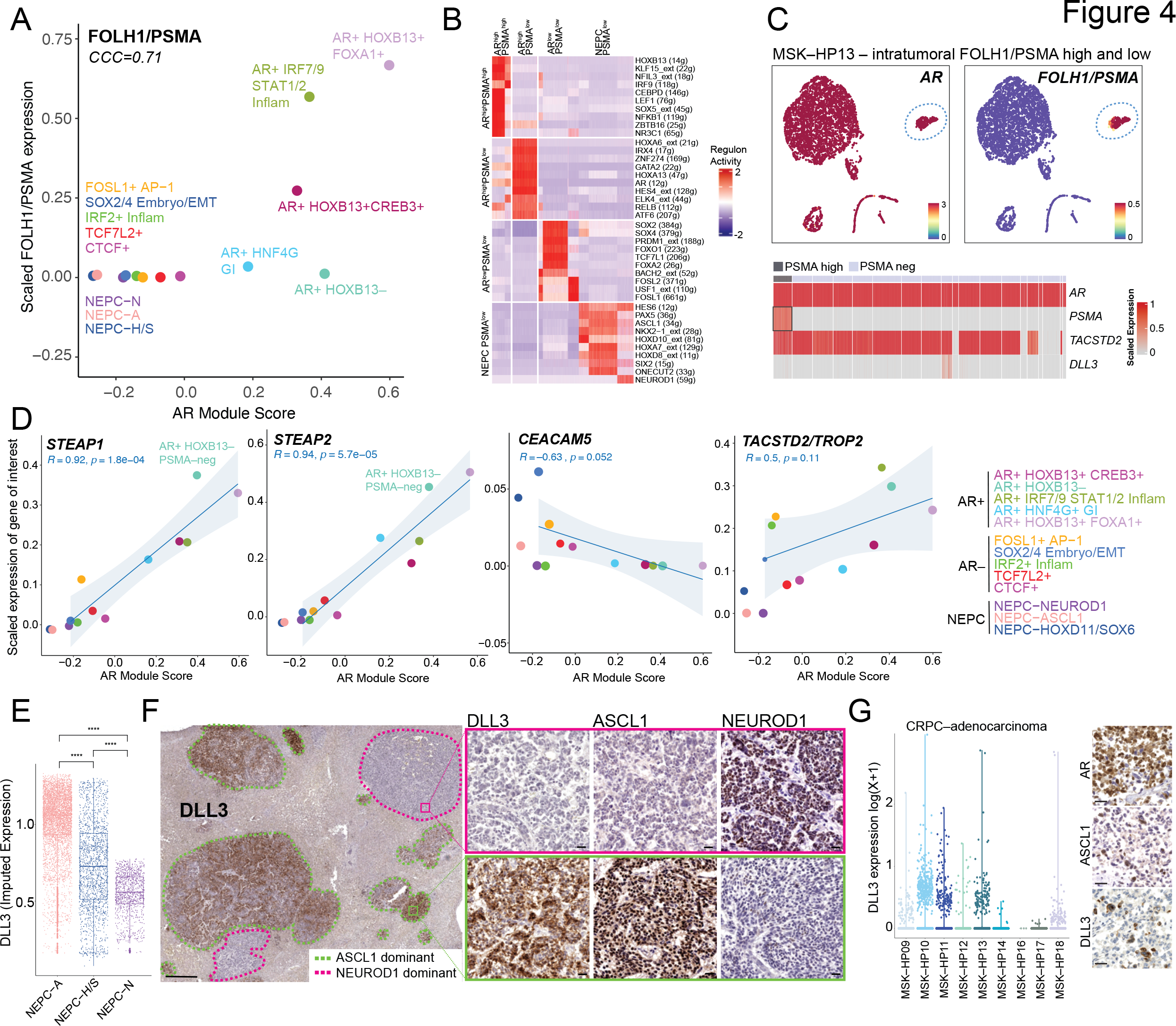
Expression of Cell Surface Markers in CRPC and NEPC GRNs. (**A**) Scatter plot with scaled FOLH1/PSMA expression (y–axis) and AR module score (x–axis) (Methods) for each GRN as colored in Figure 2D (Lin’s concordance correlation coefficient = 0.71). (**B**) Heatmap of top 10 differentially active regulons in *AR^high^FOLH1/PSMA^high^*, *AR^high^FOLH1/PSMA^low^* (from MSK– HP13), *AR^low^FOLH1/PSMA^low^*, and NEPC/*FOLH1/PSMA^low^*. *Per* cell regulon activity scores are shown (scale: –2 to 2) (Methods). (**C**) UMAP of *AR* and *FOLH1/PSMA* expression in tumor cells of MSK–HP13. Dotted circles denote region of *FOLH2*–positivity in otherwise largely *FOLH1*– negative MSK–HP13 biopsy. Heatmap of scaled expression (scale 0 to 1) is shown below with a blue box marking *FOLH1/PSMA*–positive cell population. (**D**) Scatter plots are shown of scaled expression of respective cell surface antigen (*STEAP1*, *STEAP2*, *CEACAM5*, and *TACSTD2/TROP2*, y–axis) and *AR* module score (x–axis) with each dot representing a GRN. Colors of GRNs correspond to GRN annotation on right separated by *AR*–positive, AR–negative and NEPC groups. Linear fit was calculated between two markers for *only* CRPC–adeno GRNs; the corresponding Pearson’s correlation is noted *only* for CRPC–adenocarcinoma GRNs or AR– positive and AR–negative GRNs alone. (**E**) A boxplot for *DLL3* imputed expression (MAGIC, *k=20, t=1)* is shown for NEPC–A, NEPC–H/S and NEPC–N regulons. Significance was assessed by Wilcoxon–signed rank test. Abbreviations: ****(P<1x10^-4^). (**F**) Immunohistochemistry of a liver with multiple metastases (PMID 3459916) shows distinct ASCL1–dominant (green dotted line) and NEUROD1–dominant (pink dotted line) foci prospectively stained for DLL3 expression. Zoomed images of two regions with DLL3+ and DLL3–negative foci are shown for DLL3, ASCL1, and NEUROD1 expression. Scale bar is 50 µM. (**G**) Dot plot of *DLL3* expression [non–imputed, log(X+1)] in CRPC tumor biopsies in single cell human RNA–sequencing data. This analysis suggests that a subset of CRPC adenocarcinoma cells are *DLL3* expressors. On the right, representative immunohistochemistry is shown of a biopsy with interspersed ASCL1/DLL3 cells among AR positive cells (Patient 4). Scale bar is 50 µM.

We next studied *STEAP1* and *STEAP2*, both of which showed a positive correlation with AR signaling in CRPC samples (*R*=0.92, *P*=1.8x10^-4^ and *R*=0.94, *P*=5.7x10^-5^, respectively). Of note, the *AR*+ *HOXB13*–negative GRN in the *AR*^high^*/PSMA*^low^ sample showed robust *STEAP1* and *STEAP2* expression, suggesting that co–targeting of STEAP and PSMA in AR–positive disease may be an effective strategy to achieve broader tumor cell coverage (*4*) (**Figure 4D**). In this context, we also unbiasedly identified other cell surface markers within our GRNs that could be utilized in combination with known cell surface antigens, such as *PSMA* (e.g. *CEACAM5, FGFR1, PMEPA1,* and others in **Supplementary** Figure 10D).

Turning next to *TACSTD2* (*TROP2)*, the target of a clinically approved ADC for triple negative breast and bladder cancers (with additional clinical trials underway in lung adenocarcinoma and prostate cancer), we noted *TROP2* expression in most CRPC– adenocarcinoma clusters but with no correlation with *AR* expression (**Figure 4D**). This finding is consistent with our immunohistochemistry data where TROP2 was expressed in all ADCs and nearly all HCGs (**Figure 1H**) (*37*). In contrast to *TROP2*, *CEACAM5* displayed a negative correlation with *AR* in CRPC–adenocarcinoma and was expressed in all NEPC clusters, suggesting that *CEACAM5* is an actionable target for non–AR–driven disease (**Figure 4D**) (*46*).

Given that *DLL3* is a therapeutic target for both SCLC and NEPC and is downstream of *ASCL1* (*47*), we next explored the expression of *DLL3* in our NEPC regulons. While *DLL3* was expressed in all NEPC regulons, NEPC–N regulons displayed lower expression compared with NEPC–A (**Figure 4E, Supplementary** Figure 10E–F). As there were no NEPC–N tumors represented in our TMA cohort, which consists of punch biopsies from tumor blocks (**Figure 1**), we further studied full face sections of a liver metastasis from a patient with ASCL1–positive NEPC, a case previously identified in a study of neuroendocrine chromatin landscapes (*31*). IHC analysis revealed divergent ASCL1 and NEUROD1 expression in discrete tumor foci. Whereas DLL3 was abundantly expressed in the ASCL1–positive foci, we observed little or no expression in the NEUROD1–positive foci (**Figure 4F**). While this spatial analysis is from a single patient, the collective single cell sequencing data reveal differing levels of *DLL3* expression across the NEPC spectrum. This heterogeneity could become an important variable in interpreting clinical response data in NEPC patients receiving DLL3–targeted therapy.

Because NEPC typically arises as a consequence of lineage transformation from PRAD, we next looked at *DLL3* expression in our adenocarcinoma cohort and found clear evidence of DLL3 expression within subsets of cells within CRPC–adenocarcinoma tumors (**Figure 4G, Supplementary** Figure 10G). Within these CRPC PRAD samples, a subset of the *DLL3* expressors were also positive for *CHGB* and *ASCL1* expression and scored highly for the NEPC gene signature (**Supplementary** Figure 10H). Similarly, the TMA analysis identified rare HCG tumors with mixed lineage marker expression (AR and ASCL1) that also expressed DLL3 (example shown in **Figure 4G**). Collectively, these data raise the possibility of early therapeutic targeting of rare NEPC cells in tumors with high–grade morphology or plasticity–associated genotypes, such as *TP53* and/or *RB1* loss.

## DISCUSSION

Multiple cancer types, after treatment with next generation targeted inhibitors, can evolve to develop an array of heterogenous lineage states––a process often referred to as lineage plasticity (*9*). Prostate cancer serves as an archetype for the emergence of such plastic drug– resistant cell states, typified by the transformation from adenocarcinoma to neuroendocrine cancer (*10*). These cell states in prostate cancer as well as other tumor types are generally associated with poor responses to signaling inhibitors, current cell surface–based therapeutics or chemotherapeutics (*11*). While there has been growing insight into the different cell states that may emerge in both mouse and human prostate tumors, an understanding of the diversity of transcriptional networks underlying these cell state changes and their associated lineage and cell surface marker expression in plastic prostate tumors remains limited.

To enhance our understanding of these lineage states and how they relate to cell surface marker expression, we pursued two parallel approaches: (i) annotation of individual marker gene expression across an extensive TMA panel of late–stage PRAD and NEPC samples, and (ii) single cell transcriptome analysis to identify distinct cell states, as well as the putative GRNs associated with those states, with subsequent linkage back to individual marker gene expression. The single marker gene approach confirmed that both YAP1 and TROP2 are robust markers for PRAD and HGC histology, whereas DLL3 is an exclusively NEPC–specific cell surface antigen. However, deeper analysis of the NEPC state with additional markers (SYP, ASCL1, NEUROD1 and INSM1) revealed important caveats that could help refine and sharpen the clinical diagnosis of NEPC. For example, both ASCL1 and INSM1 are highly specific markers of NEPC and tend to lose expression of YAP1 expression, providing a strong dichotomy between PRAD and NEPC states. SYP is robustly expressed in the majority of NEPC as well (and is correlated with ASCL1 expression) but is also robustly expressed in a subset of AR–positive PRAD and HGCs that <u>do not</u> express ASCL1 or INSM1 (often referred to as “amphicrine”). Thus, SYP expression alone is neither sufficiently sensitive nor specific for defining bona fide NEPC states.

Our analysis of NEUROD1 expression through TMA–based protein expression and scRNA–seq analysis is similarly revealing due to the rarity of NEUROD1–positive *versus* ASCL1– positive NEPC, particularly considering the relative frequencies of NEUROD1– *versus* ASCL1– positive SCLC. These differences could simply be a consequence of tissue/cell of origin (e.g., prostate adenocarcinoma cells *versus* lung neuroendocrine cells). However, recent studies in SCLC have established that NEUROD1–positive clones can emerge from ASCL1–positive cells, particularly in response to bottlenecks imposed by selective pressure from chemotherapy (*48, 49*). This plasticity between ASCL1– and NEUROD1–positive states may explain our detection of both populations by scRNA–seq in two NEPC samples. The fact that DLL3 expression is significantly lower in NEUROD1–positive NEPC cells may have implications for the clinical success of DLL3–targeted therapies. Taken together, these insights on *ASCL1 versus* NEUROD1 expression within NEPC and SYP expression in PRAD argue for a standardized IHC panel–based approach using ASCL1, NEUROD1, INSM1 and SYP, in conjunction with histomorphological assessment, to add greater precision to the diagnosis and treatment of NEPC.

While IHC panel–based approaches may yield improved insight into cell states, our study has demonstrated they do not capture the heterogeneity of late–stage prostate cancer, both across patients, but equally important within a single patient. Our analysis of single cell transcriptomes has thus provided further insight into the heterogeneity of cell states that underlie PRAD and NEPC. First, we observed markedly increased transcriptional diversification in CRPC and NEPC when compared to naive/CSPC tumors. This increase may be a consequence of the expanded number of putative GRNs. In addition to well–established AR–positive state (*HOXB13+* and *FOXA1*+ GRNs), we identified inflammatory and GI lineage states (*10, 43*) that have previously been implicated in ARSI resistance. The *AR*–negative GRNs included epithelial– mesenchymal and embryonic/stem (*TWIST2*, *SOX2/4*, *FOSL1/2*), inflammatory (*STAT1/2*), and WNT signaling (*TCF7L2)*. Many of these patient–derived transcriptional states are also present in GEMMs as well as human tumoroids (*10, 21*), providing further validation of the clinical relevance of these models for preclinical studies. We also note specific populations unique to murine models (e.g., *Pou2f3+* cells in the setting of prostate specific *Trp53, Rb1, Pten* deletion) that we failed to detect in our human TMA or single cell data.

We analyzed how the expression of common cell antigens used for antibody–drug conjugates, T–cell engagers, and theranostics varies as a function of transcriptional states. As expected, *AR*–positive GRNs were correlated with *PSMA* expression, with a clear exception in MSK–HP13 which demonstrated an *AR*–positive, *HOXB13*–negative regulon lacking *PSMA* expression. The latter is consistent with recent evidence implicating HOXB13 as a direct regulator of *PSMA* (*42*). However, because comparable levels of *HOXB13* activity can be present in PSMA+ and PSMA– cells (for example, see MSK–HP13, **Supplementary** Figure 7), HOXB13 expression alone is not sufficient. Towards this, *AR*^high^*/PSMA*^high^ and *AR*^high^*/PSMA*^low^ networks could be useful in identifying these additional *PSMA* regulators. Another clinically relevant finding driven by our analysis is the tight correlation of *STEAP1* and *STEAP2* expression with all AR^high^ regulons regardless of *PSMA* status, raising the potential for STEAP–targeted ADC therapy alone or in combination with PSMA–directed RLT (*4*). This is further supported by immunohistochemical analysis of the rapid autopsy tissues, which demonstrated a lower proportion of PSMA–high (45%) *versus* STEAP1–high (70%) tumors (*50*). *TROP2* expression was enriched across all adenocarcinomas regardless of AR status, indicative of a broader lineage profile (*37*).

In summary, our findings provide a comprehensive atlas of progressive heterogeneity in late–stage prostate cancer, including the identification of putative transcriptional networks and their association with lineage and cell surface markers. As far as potential limitations (site of biopsy, time of processing, batch correction, etc.) are concerned, identification of such GRNs require additional validation in larger cohorts, ideally linked with chromatin accessibility data (e.g., multiome). With recent advances in comprehensive molecular diagnostic liquid assays, one can envision incorporation of GRN–based classification as an additional tool to refine patient selection and therapy decisions.

## Data and Material Availability

Human raw data for a subset of Naïve and CSPC samples, as per *Karthaus et al* (*Science,* PMID 32355025) are available at the Data Use and Oversight System controlled access repository https://duos.broadinstitute.org/ [accession no. DUOS-000115, samples: HP95 (MSK–HP01), HP96 (MSK–HP02), HP97 (MSK–HP03), HP99 (MSK–HP04), HP100 (MSK–HP05), and HP101 (MSK–HP06)]. Human raw data and 10X formatted files for CRPC samples are available at Gene Expression Omnibus repository [GSE210358, Chan*, Zaidi* et al, *Science*, PMID 35981096). For previously unpublished samples, HMP22 (MSK–HP16), HMP23A/B (MSK–HP07), HMP24 (MSK–HP08), FASTQ and 10X files have been upload to Gene Expression Omnibus repository, along with two RDS files that contain all and tumor cells, respectively (accession ID pending). GEMM raw data and 10X formatted files for WT, PtR and PtRP are available at Gene Expression Omnibus repository [GSE210358, Chan*, Zaidi* *et al*. *Science*, PMID 35981096]. Code for notebooks to reproduce figures will be available at GitHub (in process). All other data are available in the manuscript or the supplementary materials.

## Acknowledgments

Authors are incredibly grateful to the prostate cancer patients who participated in this research. We are also grateful to the patients and their families, Celestia Higano, Evan Yu, Elahe Mostaghel, Heather Cheng, Michael Schweizer, Jessica Hawley, Bruce Montgomery, Andrew Hsieh, Jonathan Wright, Daniel Lin, Funda Vakar-Lopez, and the rapid autopsy teams for their contributions to the University of Washington Prostate Cancer Donor Rapid Autopsy Program. We further appreciate the efforts of the Memorial Sloan Kettering Cancer Center (MSK) Genitourinary faculty for recruitment of prostate cancer tumor specimens. We thank the MSK core facilities for their invaluable help, namely the Molecular Cytology Core and Pathology Core for their help with confocal microscopy and IHC. This work is supported by R01 CA234715; R01 CA266452-01; R21CA277368-01; P50CA097186; the Institute for Prostate Cancer Research and the Prostate Cancer Foundation. S.Z. is supported Prostate Cancer Foundation Young Investigator Award, Louis V. Gerstner Jr. Scholarship, NIH K08 CA282978, and Burroughs Wellcome Fund Career Award for Medical Scientists. J.C. is supported by the National Research Foundation of Korea (NRF2022R1A4A2000827). P.S.N. is supported by NCI P50CA097186, U54CA224079, R01CA234715 and Challenge Awards from Prostate Cancer Foundation. M.C.H is supported by the Department of Defense (DoD) Prostate Cancer Research Program (W81XWH-21-1-0229, W81XWH-20-1-0111) and by Grant 2021184 from the Doris Duke Charitable Foundation. C.L.S. is supported by HHMI; National Institute of Health (CA193837, CA092629, CA224079, CA155169, CA265768, and CA008748), and Starr Cancer Consortium (I12–0007).

## Author contributions

S.Z., M.H., and C.L.S. conceived the project. S.Z., M.H., and C.L.S wrote the manuscript. H.I.S, D.E.R, and M.J.M provided human tumor specimens for scRNA–seq. A.G., interpreted histology and staining for prostate cancer patients for scRNA–seq. S.Z., J.L.Z., W.R.K and P.A.W. performed human tumor tissue dissociation. P.S.N., C.M. and M.P.R. enrolled patients into rapid autopsy protocols and provided rapid autopsy biospecimens. K.M.W and D.G. provided GEMMs for scRNA–seq. M.P.R., A.B. A.J., N.R., J.S, L.D.T, R.A.P, E.S, and M.C.H. performed stains, interpreted and/or scored human tissue microarray. O.C., I.M., and T.X. performed single-cell sequencing. L.M. and R.C. oversaw the single-cell–sequencing experiments. S.Z. J.P, D.H.K. A.O, J.C., and J.C. performed computational analyses. M.H. and C.L.S. oversaw the project.

