## Supplementary Information for "Single Cell Analysis of Treatment–Resistant Prostate Cancer: Implications of Cell State Changes for Cell Surface Antigen Targeted Therapies"

**This PDF file includes:**

Materials and Methods

Supplementary Figures 1–10

Supplementary Figure Legends

Supplementary Tables 1–16 (separate attachments)

#### METHODS

##### Human Tissue Microarray

**Human tissue microarray.** Tissue samples were procured from men who died of metastatic CRPC–adenocarcinoma and who signed written informed consent to undergo a rapid autopsy as part of the Prostate Cancer Donor Program at the University of Washington as described previously (*1-4*). The Institutional Review Boards of the University of Washington and of the Fred Hutchinson Cancer Center approved this project. In this study, a total of 131 tumors including 42 adenocarcinomas (PRAD), 51 high–grade carcinomas (HGC), and 38 high–grade neuroendocrine prostate carcinomas (NEPC) from 16 patients were analyzed (**Figure 1, Supplementary Figure 1A** and **1C**). PRAD was defined as a carcinoma with discernable gland formation, ranging from small individual glands to large cribriform glands with nuclei showing mostly open chromatin and prominent nucleoli. HGC was defined as a poorly differentiated carcinoma without gland formation, indistinguishable from a poorly differentiated carcinoma arising in another organ site. NEPC was defined as a carcinoma with small cell features (characteristic “small blue cells”) with minimum cytoplasm, indistinct cell borders, and lack of prominent nucleoli. TMA containing 2 separate cores for each metastatic site was constructed and adjacent sections were stained with antibodies specific to a total of 13 proteins including AR, NKX3.1, CK8, TP63, YAP1, SYP, INSM1, ASCL1, NEUROD1, POU2F3, MYC, FOXA2, SOX2, EZH2, TFF3, TROP2, DLL3, KI67 (antibody specifications listed in **Supplementary Table 1**). Positive controls for NEUROD1, P63, and POU2F3 are shown in **Supplementary Figure 1B**, as these genes demonstrated negative staining in most of our samples. Ischemic time shown in **Supplementary Table 2** was defined as the interval between the time of death to time of necropsy. Pearson’s correlation was calculated between the mean of H–scores for each protein across all tumors from a given patient *versus* ischemia time. Tissue sections were counterstained with hematoxylin and slides were digitized on a Ventana DP 200 Slide Scanner (Roche). All cases were reviewed by experienced genitourinary pathologists (M.R., M.C.H.) and immunoreactivities were scored in a blinded manner using a previously established H–score system, whereby the optical density level (“0” for no brown color, “1” for faint and fine brown chromogen deposition, and “2” for prominent chromogen deposition) was multiplied by the percentage of cells at each staining level, resulting in a total H–score (range 0–200). The final H–score for each sample was the average of two replicate tissue cores. For KI67, any nuclear positivity was counted, and the percent of positive nuclei is shown (range 0–100). Included patient identifiers and H–scores for each protein are detailed in **Supplementary Table 1**.

**Comparison H–Score Trends for Lineage Markers Across PRAD/HGC/NEPC**. For each histology (PRAD, HGC, and NEPC), H–scores *per* marker gene were compared using a modified Wilcoxon–ranked test (please refer to https://rdrr.io/github/chvlyl/ZIR/man/ziw.html). P–values were adjusted using Bonferroni correction for the number of tests within each gene (**Figure 1B–D** and **H–I**, **Supplementary Figure 1A**). For YAP1 and EZH2, H–scores were also compared between NEPC *versus* PRAD plus HGC or PRAD *versus* HGC plus NEPC respectively, using a modified Wilcoxon–ranked test. Furthermore, scatter plots were generated for YAP1, SYP, DLL3, INSM1, or TROP2 *versus* ASCL1 or NEPC score (defined as mean H-score of ASCL1, INSM1, and FOXA2), or EZH2 *versus* Ki67. To evaluate for concordance between the H–scores of two genes of interest, Lin’s concordance correlation coefficient (CCC) was calculated (**Figure 1F–1G** and **Supplementary Figure 1D** and **1F**)**.** For genes without clear linearity, CCC was not calculated, but rather the data was simply visulazed and displayed as a scatter plot (**Supplementary Figure 1C** and **1E)**. For *EZH2* and *Ki67*, as the H–score and proliferative index were on different scales (either 0–200 or 0–100, respectively), Pearson's correlation was used to define the strength of correlation (**Figure 1E**).

#### Sample Handing and Characteristics for Single Cell RNA–Sequencing

**Patient derived tissue.** Tumor tissues were collected from patients undergoing a surgical resection or tissue biopsy for clinical care at Memorial Sloan Kettering Cancer Center (MSKCC). Informed consent was obtained for all patients and approved by MSKCC's Institutional Review Board (IRB) #12–245 (NCT: 01775072), #06–107, and #12–001. Tumor biopsies included 9 naïve or castrate–sensitive prostate cancer specimens (8 patients, 2 specimens were obtained from one patient: MSK–HP07A/B, Naïve or CSPC) and 14 metastatic castrate–resistant prostate cancer specimens (11 CRPC–adenocarcinoma and 3 NEPC specimens, from 13 patients, 2 specimens were obtained from one patient: MSKC–HP11A/B). Of note, 6 out of the 9 naïve/CSPC biopsies were previously reported by our group (*5*) and are available at the Data Use and Oversight System controlled access repository: https://duos.broadinstitute.org/ (accession no. DUOS–000115, files annotated as HP95T to HP101T or renamed as MSK–HP01 to HP06, all FASTQ files were re–processed '**Preprocessing of scRNA–sequencing dataset**'). All CRPC biopsies were collected prospectively from 2019 to 2022. An initial analysis of 13 out of 14 biopsies was previously reported specifically in the context of JAK/STAT signaling (*6*).

Samples were freshly derived from tumor sites from patients with untreated (naïve) or CSPC, or CRPC. All CRPC patients in this cohort had progressed after treatment with antiandrogen therapy and a next–generation anti–androgen therapy (ARSI), with or without a taxane. Serum PSA levels were reported from the date one week before or after of biopsy. Detailed clinical characteristics including AJCC staging, location, PSA, type and number of prior treatments, and genomic alterations are listed in **Supplementary Table 3**. Routine anatomic pathology and immunohistochemistry were performed by a MSKCC pathologist (A.G.). For CRPC single cell biopsies, distinct from the categorization of TMA samples, patients were classified into either CRPC adenocarcinoma (CRPC–adeno) or neuroendocrine with characteristic small cell features (NEPC).

**Targeted sequencing of human tumor tissue.** For patients who consented to IRB 12–245 (NCT 01775072; MSK–IMPACT) (*7*) and for clinical purposes, targeted sequencing was performed for a panel of actionable cancer genes for alterations (available for 14 patients, **Supplementary Table 3**, also subset previously reported in (*6*)). Genomic data was collected on retrospective review of the electronic medical record.

**Sample processing of tissue.** Fresh benign and malignant tissue was mechanically cut using a scalpel into small pieces (~1–5 $mm^3$). The tissue was then processed and dissociated in 5–10 mg/ml collagenase type II (Gibco) solution in adDMEM/F12+/+/+ with 10 µM Y–27632 dihydrochloride for 30 minutes to 2 hours on a 37$^{o}$C shaking platform. This was followed by a 1 minute 0.5 M EDTA wash at room temperature, and subsequent digestion with TrypLE (Gibco) with 10 µM Y–27632 dihydrochloride for 5–10 minutes at 37$^{o}$C on a shaking platform until a single cell suspension was obtained. If more than 10% of doublets were present (visual inspection) or there was evidence for <80% viability (hemocytometer, using 0.2% Trypan Blue), then cells were FACS–sorted for singlets and viability using a DAPI.

**Sample processing: 10X single cell RNA–sequencing.** Dissociated cells were subjected to scRNA–seq using 10X genomics Chromium Single Cell 3' Library and Gel bead Kit (v3 for human, except v2 for HP95T as previously described (*5*)) per manufacturer's protocol. ~3000 to 10,000 cells per sample was encapsulated and barcoded following the manual. Sample viability varied between 72 and 95% (0.2% Trypan Blue). The final sequencing libraries were double–size purified (0.6–0.8X) with SPRI beads and sequenced on Illumina Nova–Seq platform (R1–26 cycles, i7–8 cycles, R2–70 cycles or higher). For human samples, on average, 4,864 cells per clinical biopsy were sequencing at a depth of ~ 82,429 reads per cell (~258 million reads per sample). The unique mapping was high, between 62.9–82.6% (except MSK–HP07A/B 58.6% and 60.5%), with a median number of unique transcripts per cell being 7,492.

**Single Cell RNA–Seq Processing and Analyses**

**Preprocessing of scRNA–sequencing dataset.** For each human sample, FASTQ files produced from 10X scRNA–seq were mapped to the pre–built human reference (hg38) and converted to counts using the pipeline "cellranger count" (*8*). Cells were filtered out (or removed) based on two criteria: (1) high fraction of mitochondrial molecules > 25% and (2) low library complexity expressing very few unique genes (<200 genes). Furthermore, putative doublets were removed using the 'DoubletFinder' package (*9*). Combining samples in the entire cohort of Naïve or CSPC, CRPC–adenocarcinoma and NEPC yielded a count matrix of 119,083 cells x 33,473 transcripts with a median of 8,268 molecules per cell and a median of 4,096 cells per biopsy. The count matrix was then normalized by (1) feature counts divided by total feature counts in each cell, (2) scaled by 10,000 ('scale.factor'), and (3) subjected to natural–log transformation with a pseudo–count of 1 ('NormalizeData'). Thereafter, cell–cycle signatures were calculated ('CellCycleScoring') and the signature scores were regressed. The features were then scaled and centered ('ScaleData'). Principal component analysis (PCA) was based on 2000 highly variable genes (HVG) (identified using 'FindVariableFeatures', excluding mitochondrial, ribosomal, and with *MALAT1*, *UBA52*, *NEAT1*, *TMSB4X*, and *TMSB10*—hererin referred to house–keeping genes) and PCA analysis was performed on the scaled matrix of HVGs with the top 30 principal components (PCs) detected by knee–point (33% variance explained).

#### Batch Correction

**Batch correction for subsetted coarse cell type.** Batch correction was not necessary for the identification of coarse cell types (i.e. epithelial/neuroendocrine, lymphoid, myeloid, and stromal). However, upon subsetting each coarse cell type, patient–level batch effect was observed with cells clustering by patient rather than by cell type (e.g. multiple Treg populations with *FOXP3* expression). Therefore, batch correction was performed on each coarse cell type using fastMNN (further detailed below in **'Human Cell Type Annotation'**) due to the ability to perform hierarchical merging similar histologies (i.e. CRPC–adenocarcinoma or NEPC). To evaluate the effect of batch correction, we utilized an entropy–based measure that quantifies how much normalized expression mixes across patients (*10, 11*). This demonstrated that immune cells had the highest entropy (followed by stromal), whereas epithelial/tumor cells had the lowest entropy suggestive of increased inter–tumoral diversity (**Supplementary Figure 3G**). Furthermore, by examining the expression of specific cell type markers, we did not observe an over–correction of granular cell types (e.g. normal epithelium *versus* tumor, neuroendocrine, Tregs, macrophage and immune subsets, etc.). These results thus balanced the need for batch correction, while maintaining true biologic heterogeneity.

#### Human Cell Type Annotation

**Human coarse cell type identification.** We then used a hierarchical strategy for cell typing, initially at coarse resolution (epithelial, immune, *versus* stroma), and granular cell types (normal *versus* tumor, neuroendocrine, etc.) using canonical marker gene expression within unsupervised clusters. For coarse resolution cell typing, clustering was performed by first constructing a *KNN* graph on the Euclidean distance in PCA space with 30 principal components (PCs) as identified by knee–point. Edge weights between two cells were adjusted based on Jaccard similarity ($k$=30, based on adjusted Rand index), generating a shared nearest–neighbor graph (SNN). Louvain clustering was applied to the SNN with resolution=0.5, resulting in 43 clusters spanning 119,083 cells ('FindNeighbors' and 'FindClusters' functions). Clusters were annotated based on expression of canonical markers, including *EPCAM* or *SYP/CHGA/CHGB* for epithelial/neuroendocrine cells (23 clusters, 68,816 cells); *PTPRC* and either *CD3E*, *CD79A*, or *KLRD1* for lymphoid cells (8 clusters, 21,044 cells); *PTPRC* and *CD14* and/or *LYZ* for myeloid cells (4 clusters, 10,126 cells); and *COL1A2, ACTA2* and *PECAM1* for stromal cells (8 clusters, 19,097 cells).

**Human designation of epithelial and tumor cells.** For EPCAM–positive or SYP/CHGA–positive clusters (N=68,816 cells, including 28,508 CSPC cells, 24,866 CRPC-adenocarcinoma cells, and 15,442 NEPC cells), we projected scaled counts onto the top 30 PCs based on knee–point, restricting to 2,000 highly variable genes and a set of biologically relevant genes (AR, JAK/STAT, EMT, NEPC–related, (*6*), found in **Supplementary Table 13**), corresponding to 28.8% of variance explained. Louvain clustering (as described above) was performed with resolution of 0.5 generating 31 clusters. To identify malignant tumor cells within subsetted epithelial/neuroendocrine cells, we evaluated the expression of genes known to be upregulated in malignant prostate cancer, including *AR*, *KLK*2, *KLK3*, *FOLH1*, *STEAP1*, *STEAP2*, *TACSTD2*, *ERG*, *ETV1*, *ETV6*, *ASCL1*, *NEUROD1*, *DLL3*, *CHGA*, and *CHGB* (**Supplementary Figure 3B**) (*12*). Specifically, 20 out of 31 clusters showed robust expression of at least 2 markers. If clusters were largely derived from NEPC samples (MSK–HP19, MSK–HP20, MSK–HP21) and showed expression of *SYP*, *CHGA*, and *ASCL1* or *NEUROD1*, the clusters were labeled as NEPC. The majority of non–tumor cells (including basal, club cells, and lowly expressing luminal cells) (*5*) were derived from primary tumors, which are well known to be admixed with benign cells.

Putative tumor cells were again subsetted (N=35,962 cells), and subjected to batch correction using FastMNN, (*13*) which was applied to the normalized count matrix and reduced to the top 30 PCs (refer to section ‘**Batch correction**’ for rationale). Here, we performed Phenograph clustering (*k=40)*, which resulted in 32 clusters. All but one cluster showed consistent expression of tumor associated markers, except for cluster 6 (N=266 cells) which exhibited the distinct expression of *ALB, CRP*, *VTN*, *A1CF*, *GC*, and *AMBP*, known to be expressed in hepatocytes, and comprised of cells from patients MSK–HP19, MSK–HP09, MSK–HP17, and MSK–HP18, where the biopsy was derived from liver metastases **(Supplementary Figure 3C).** We thus concluded that this cluster comprises of benign hepatocytes, and thus was removed from the final set of tumor cells.

These malignant cells, without hepatocytes, were subsetted (N=35,696 cells; 4,001 CSPC cells, 20,088 CRPC–adenocarcinoma cells, and 11,607 NEPC cells) and underwent batch correction (refer to section ‘**Batch correction**’ for rationale). FastMNN was applied to the normalized count matrix, reduced to the top 20 PCs. Phenograph clustering (*k*=30) was performed to identify 31 clusters, all of which showed expression of at least 2 tumor associated markers (**Supplementary Figure 3B)**. As a confirmatory approach, we assessed single–cell copy number variation (CNV) profiles in tumor cells, by utilizing inferCNV package, which measures the expression of genes across chromosome positions and compares them to a pre–designated reference set of cells (myeloid immune cells). To identify CNV profiles, we used a sliding window approach of 101 genes and considered any deviation from the reference mean of at least 1.5 standard deviation as a copy number variation (using inferCNV package). As expected, we observed an increased number CNVs in malignant cells with highest burden in NEPC (*14*) consistent with prior reports and providing additional support for accurate identification of malignant tumor cells (**Supplementary Figure 3D)**. We classified each Phenograph cluster into Basal, LumA, and LumB using the pre–defined PAM50 model from genefu (https://bioconductor.org/packages/release/bioc/ html/genefu.html). Considering that our tumor cells lacked HER2–positivity and that we already filtered out normal cells, we excluded these categorizations from our PAM50 classification. (*15*)

**Human immune and stromal annotation.** Stromal, myeloid, and lymphoid cells were subsetted based on the following respective markers: (1) *COL1A2, ACTA2* and *PECAM1* for stromal cells (8 clusters, 19,097 cells); (2) *PTPRC* and either *CD3E*, *CD79A*, or *KLRD1* for lymphoid cells (8 clusters, 21,044 cells); (3) *PTPRC* and *CD14* and/or *LYZ* for myeloid cells (4 clusters, 10,126 cells). For each subsetted cell type, fastMNN was applied to the normalized count matrix, and then reduced to the top 30 PCs. Refer to '**Batch correction for subsetted coarse cell type'** for rationale.

**<u>Measuring inter–patient heterogeneity per cell type</u>**

For each tumor type, including Naïve or CSPC, CRPC–adenocarcinoma, and NEPC, we employed an entropy–based approach to assess interpatient diversity, as has been previously described (**Figure 2C** and **Supplementary Figure 3F** and **3H)** (*10, 11*). We used Phenograph clusters ($k$=30) derived from a subset of tumor cells (N=35,696 cells), myeloid, lymphoid, and stromal populations. Each cluster $C$ represents a discrete phenotype of a given cell. To account for varying cell numbers across clusters and cell types, we implemented a subsampling strategy. Specifically, we randomly selected 100 cells from each cluster, repeating this 100 times with replacement. We calculated the Shannon entropy of patient frequencies $P$ in each subsample, denoted as $H_c$:

$$H_C = \sum_P \quad -q_P \log q_P .$$

To avoid bias introduced by repetitive sampling of a limited cell pool, clusters containing fewer than 100 cells were excluded from the calculation. The distribution of Shannon entropies, bootstrapped from clusters, was analyzed, and compared using a Bonferroni-adjusted Wilcoxon signed-rank test.

#### Identification of Single–Cell Regulatory Network Inference

**Human regulatory network inference.** To understand the TF networks that may account for the observed diversity in CRPC–adenocarcinoma and NEPC tumors, we utilized single–cell regulatory network inference (SCENIC) (*16*) and obtained a regulon–activity matrix for each cell. The regulon–activity was than z–scaled as is shown in **Figure 2D** and **Supplementary Figure 4A** and **4C**. To obtain a set of distinct GRNs that contribute to the observed heterogeneity in CRPC and NEPC samples, we hierarchically clustered Phenograph clusters on the scaled regulon activity matrix, employing the Ward D2 method (Euclidean distance) for both CRPC–adenocarcinoma and NEPC clusters. To define common and distinct gene–regulatory networks (GRNs, group of regulons) across samples in an unbiased fashion, we applied a dendrogram height cutoff of 15 based on the adjusted Rand index (ARI) of CRPC–adenocarcinoma (overall >= 0.65 indicating moderate to high recovery) (**Supplementary Figure 4B**), and for consistency, we applied the same threshold for NEPC. This yielded 10 and 5 GRNs for CRPC–adenocarcinoma and NEPC, respectively. Among NEPC GRNs, the 2 smallest GRNs with cells < 600 (NEPC–2 and NEPC–3) lacked distinct regulon activity from their nearest neighbor, and therefore were merged (NEPC–2 with NEPC–HOXD11/SOX6 GRN and NEPC–3 with NEPC–A GRN). This resulted in 10 and 3 final GRNs for CRPC–adenocarcinoma and NEPC, respectively. To determine the order of 13 defined GRNs, we additionally performed hierarchical clustering on each set of the 10 CRPC–adenocarcinoma or 3–NEPC GRNs, generating the dendrogram shown in the heatmap in **Figure 2D.** For each GRN, we calculated regulon specificity scores (RSS) (*17*) on the raw regulon activity matrix with the function calcRSS in SCENIC based on Jensen–Shannon divergence. We then z–scaled the RSS, assigning each regulon to the GRN group where it scored the highest. To rank regulons in each GRN by their activity, we further performed Student's t–test comparing the mean value of a regulon in a given GRN versus all other GRNs. Significantly active regulons within each GRN were defined using the criteria of a p–value < 0.01, a mean regulon activity value ≥ 1.0, and a mean difference of regulon activity in a given GRN *versus* all other GRNS ≥ 1.0. The mean value and regulon ranking within each GRN can be found in **Supplementary Tables 5** and **6**, respectively.

**Robustness Analysis of Human GRNs.**

**A. Subsampling cells:** We iteratively subsampled cells to ensure the robustness of our GRN analysis. We randomly sampled 10, 20, 50 and 100 cells from each Phenograph cluster and then performed an identical set of downstream analyses as with our initial/complete dataset. This includes the following: (1) inferring the regulon activity with SCENIC (*16*), (2) z–scaling of the regulon activity, (3) hierarchical clustering of Phenograph clusters (*k=20)* for both CRPC–adeno and NEPC, (3) selection of the h–cutoff based on ARI and (4) subsequent GRN grouping (h=12 for 10 cells random sampling yielding 15 CRPC GRN and 7 NEPC GRN, h=13 for 20 cells random sampling yielding 15 CRPC GRN and 8 NEPC GRN, h=13 for 50 cells random sampling yielding 14 CRPC GRN and 8 NEPC GRN, and h=12 for 100 cells random sampling yielding 14 CRPC GRN and 8 NEPC GRN), (5) order GRNs by additional hierarchical clustering, and (6) calculation of RSS. For additional details on the complete run, please refer to '**Human regulatory network inference.**' To examine the reproducibility of GRN groups with regulon activity, we calculated the ARI between down–sampled GRN groups and our complete dataset with defined human GRN using all cells. All datasets showed moderate recovery (ARI for 10 cells random sampling = 0.69, ARI for 20 cells random sampling = 0.75, ARI for 50 cells random sampling = 0.84, ARI for 100 cells random sampling = 0.75). A Venn diagram also showed the number of overlapping TFs between then down–sampled results and all cells with the significance of the overlap assessed using Fisher's exact test. All down-sampled datasets exhibited a significant overlap of TFs (Fisher's exact test p-value < 0.05) (**Supplementary Figure 5A** and **5B**).

**B. Subsampling patients:** To further ensure the robustness of our GRN analysis, we subsampled 100 cells in 6, 8, and 10 CRPC or NEPC patients (of 13 total CRPC and NEPC patients) and repeated this process iteratively 10 times with replacement. For each subsampling run, we utilized SCENIC to identify enriched TF/regulons and calculated the proportion of overlapping TFs/regulons between our initial GRNs (**Figure 2D**) and newly identified subsampled GRNs. For initial and subsampled GRNs that share more than 50% of TFs/regulons, we utilized a Fischer' exact test to determine whether the overlap was significant: *P–value* < 0.05—constituting a shared and recurrent GRN. By subsampling 6, 8, and 10 patients, we identified eight, nine and nine recurrently identified GRNs that significantly overlapped in our initial GRN assignments (and were found in more than 5 of the iterations) (**Supplementary Figure 6**). Several of the recurrently noted GRNs had intact *AR* expression (*AR*+ *CREB3*+, *IRF2*+ Inflammatory, *AR*+*HNF4G*+, *AR*+*HOXB13*+*FOXA1*+). Of note, *AR*+ clusters also demonstrated the most mixing between tumors (as defined by patient entropy, **Supplementary Figure 2F**). We also noted recurrent GRNs for non–*AR*–driven pathways, including *TCF7L2*+, *FOSL1*+ *AP-1*, *SOX2/4*+ Embryonic EMT, and the NEPC GRNS NEPC–A, NEPC–N, and NEPC–H/S.

**GEMM regulatory network inference** For the GEMM analysis, we obtained previously processed and published scRNA–seq data (*6*) for which we similarly obtained a regulon–activity matrix for each cell, and z–scaled the matrix. As above, hierarchical clustering was performed on Phenograph clusters (*k*=30) using the Ward2 method (Euclidean distance) on a scaled regulon activity matrix, and the GRN groups were defined based on a cutoff of h=12 (ARI overall >= 0.64, the second stabilized cutoff) (**Supplementary Figure 7**). Any GRN less than 400 cells, which by visual inspection lacked distinct groups of unique regulons, was merged to its nearest neighbor.

This resulted in 9 final GRNs (**Figure 3A**). To further determine the order of the defined GRNs, we additionally hierarchically clustered the 9 GRNs, generating the dendrogram shown in the heatmap in **Figure 3A**. For each GRN, we then calculated regulon specificity scores (RSS) on the raw regulon activity matrix with the function calcRSS based on Jensen–Shannon divergence in SCENIC (*17*). We then z–scored the RSS, assigning each regulon to the GRN group where it scored the highest. To rank regulons in each GRN by their activity, we further performed Student's t–test comparing the mean value of a regulon in a given GRN versus all other GRNs. Significantly active regulons within each GRN were defined using the criteria of a p–value < 0.01, a mean regulon activity value ≥ 1.0, and a mean difference of regulon activity in a given GRN *versus* all other GRNS ≥ 1.0. The mean value and regulon ranking within each GRN can be found in **Supplementary Tables 8** and **9**, respectively.

**Robustness Analysis of GEMM GRNs.** Similar to the human SCENIC robustness analysis, 100 cells were randomly sampled from each Phenograph clusters and were subjected to an identical set of downstream analyses as with our initial/complete dataset. ARI was calculated between down–sampled GRN groups and our complete dataset GRN groups. We noted moderate recovery with an ARI = 0.74. Furthermore, the Venn diagram was constructed to show the number of overlapping TFs between the down–sampled and complete dataset (Fisher's exact test p-value < 0.05) (**Supplementary Figure 7C**).

<u>**Differential Expression and Pathway Analysis**</u>:

**Identifying DEGs and MAST.** For all differential expression, we used the 'FindAllMarkers' function, implementing MAST *v.*1.22.0. This package provides a flexible framework for fitting a hierarchical generalized linear model to the expression data. A regression model was executed as below:

$$Y_{i,j} \sim \text{condition}$$

The condition represents the condition of interest and $Y_i$ is the expression level of gene *i* in cells of cluster *j*, transformed by natural logarithm with a pseudo–count of 1. Significantly differentially expressed genes were considered with an adjusted p–value < 0.05 and absolute $\log_2$fold–change ($\log_2$FC) > 0.4 (*18*).

**GRN pathway annotation.** For each GRN, we conducted two complementary approaches: (1) gene set enrichment analysis (GSEA) using the fGSEA package, and (2) over–representation analysis (ORA) with the clusterProfiler package (*19*) on a curated set of genes (*6*) (**Supplementary Table 15** – gmt file). GSEA was performed on pre–ranked genes using 10,000 permutations. Gene ranks were calculated based on MAST differential expression result applying the following formula (Wald test statistics formula):

$$stats = qnorm(p\_val/2, lower.tail = F) * sign(avg\_log2FC)$$

For genes with an infinite value of statistics due to a p–value equal to 0 or 1, the statistics was replaced with the highest or lowest value of the statistical metric. ORA was performed on top 5 TFs based on RSS scores and their associated target genes for each GRN (**Supplementary Table 14** – list of top 5 TFs and target genes) (*20*).

**Comparison of Human and GEMM GRNS.** To compare human and GEMM GRNs, we implemented both GSEA and over–representation analysis (ORA). These analyses identified which human GRNs were enriched/overrepresented in GEMM GRNs and *vice versa*. We curated the top 5 TFs based on RSS scores and their target genes as noted by SCENIC, to create gene sets for both Human and GEMM GRNs (**Supplementary Table 14**). For inter–species comparison, we mapped mouse genes to their human counterparts (and *vice versa*) using the biomaRt package (https://bioconductor.org/packages/release/bioc/html/biomaRt.html), and discarded genes without orthologs.

We then performed differential expression analysis (refer to **'Identifying DEGs and MAST'**) to generate the input for GSEA and ORA analyses. GSEA was performed for each GRN. The genes were pre–ranked using the statistics derived from the results of the MAST differential expression analysis (refer to **'GRN pathway annotation**'). We then used 10,000 permutations to assess gene set enrichment. Simultaneously, ORA was performed for each GRN on differentially expressed genes *per* MAST with the cutoff of average $\log_2$FC > 0.25 and adjusted p–value < 0.05 on gene set in **Supplementary Table 14**. In the GSEA approach, the normalized enrichment score (NES) served as our key index to assess the enrichment between DEGs and gene lists (TFs and their regulons). For ORA, we derived an equivalent measure by computing the observed–expected ratio:

$$Obs/Exp = Gene\ Ratio/Background\ Ratio$$

Where the gene ratio and background ratio were computed as:

$$Gene\ Ratio = \#\ genes\ overlapped\ with\ specific\ category\ /\#\ genes\ overlapped\ with\ whole\ geneset$$

$$Background\ \ Ratio = \#\ genes\ in\ specific\ category/\#\ genes\ in\ whole\ geneset$$

To identify similar GRNs between human and GEMM GRNs, we select pairs of human–GEMM GRNs that displayed both a significant enrichment through GSEA and ORA (adjusted p–value < 0.05) in both comparative directions (GEMM *versus* human, and *vice versa*) (**Supplementary Figure 8**). To further prioritize TFs leading to the observed similarity, we specifically chose human–GEMM GRN pairs where at least one of the 5 top transcription factors was shared.

**Comparison of Human and GEMM with SCLC.** We performed GSEA analysis on processed data from Chan *et al*, 2021 *Cancer Cell* where subtypes were defined as SCLC–A, SCLC–N, and SCLC–P (*10*) . We assessed for the enrichment of human prostate GRNs in SCLC–A and SCLC–N, and for GEMM GRNs in SCLC–P (given that no Pou2f3 subset was found in our human dataset). MAST differential expression analysis was performed (refer to **'Differential Expression and Pathway Analysis'**) between SCLC subtypes, comparing SCLC–A *versus* SCLC–N and SCLC–P *versus* rest. Similar DEGs were determined for NEPC–A *versus* NEPC–N and GEMM–Pou2f3 *versus* rest. Scatter plots were generated with the average $\log_2$FC of SCLC–A and SCLC–N (y–axis) and NEPC–A and NEPC–N (axis) **(Figure 3C).** Genes that were significantly up–regulated (average $\log_2$FC > 0.4, adjusted p–value < 0.05) or down-regulated (average $\log_2$FC < –0.4, adjusted p–value < 0.05) were noted, with those encoding transcription factors colored in red or blue, respectively. Similar plots were generated for the Pou2f3 subsets, with transcription factors colored in brown (SCLC–P versus rest on y–axis and GEMM–Pou2f3+ versus rest on x–axis) (**Supplementary Figure 9C**).

Venn diagrams were constructed by taking the top DEGs (average $\log_2$FC > 0.4, adjusted p–value < 0.05) between NEPC–A (*versus* NEPC–N) and SCLC–A (*versus* SCLC–N) (**Figure 3D**), and GEMM–Pou2f3 (*versus* rest) and SCLC–P (*versus* rest) (**Supplementary Figure 9D**). A Fisher's exact test was used to determine significance of overlap.

#### Targeting GRN–Informed Lineage Plasticity States

**Correlation analysis of GRNs and cell surface expression.** We computed AR and NEPC module scores for each cell using published gene sets (**Supplementary Table 13**) and by implementing the 'AddModuleScore' function. The expression levels of drug target genes in each cell were standardized using min–max scaling with the lowest value constrained to 0:

$$scaled\ expression\ of\ gene\ A = [x - min(gene\ A)]/[max(gene\ A)\ - min(geneA)]$$

where the expression of gene A for a given cell is denoted by x with min(gene A) as the minimum value of all expressions of gene A, and max(gene A) as the maximum value of all expressions of gene A. The average module score or gene expression was calculated per GRN. Scatter plots were generated with Lin's concordance correlation or Pearson's correlation noted between drug target gene expression (*PSMA, STEAP1/2*, *TROP2*, *CEACAM5*, and *DLL3*) and AR module score with GRNs colored (**Figure 4A** and **4D**).

**Comparison of PSMA high and low regulons.** For PSMA expression, GRN groups were further separated into four groups ($PSMA^{high}/AR^{high}$ $PSMA^{low}/AR^{high}$ $PSMA^{low}/AR^{low}$ and NEPC) based on the scaled expression (refer to '**Correlation analysis of GRNs and cell surface expression'**) of PSMA ($PSMA^{high}$ if ≥ than median of 0.2729) and AR module score ($AR^{high}$ if ≥ than median of 0.3651). These criteria assigned 3 GRNs in $PSMA^{high}/AR^{high}$ (AR+ HOXB13+ FOXA1+, AR+ IRF7/9 STAT1/2 Inflammatory, and AR+ HOXB13+), 1 GRN in $PSMA^{low}/AR^{high}$ (AR+ HOXB13–), and 5 GRNs in $PSMA^{low}/AR^{low}$ (FOSL1 AP-1, SOX2/4 Embryonic EMT, IRF2 Inflammatory, TCF7L2 WNT, MAFG) and 3 NEPC GRNs (NEPC–A, NEPC–A/SOX6, and NEPC–N). To assess for enriched pathways, we used a previously curated sets of prostate–specific pathways, developmental pathways, and mouse cell type gene sets (67 gene sets), as well as KEGG, REACTOME, and HALLMARK canonical pathways in MSigDB v 7.1 (**Supplementary Table 15**). To identify differentially regulated regulons in four groups, we calculated regulon specificity scores (RSS) on the raw regulon activity matrix with the function calcRSS in SCENIC. We then z–scaled the RSS, ranked the regulons based on these scores and selected the top 10 TF regulons for each group. A heatmap was generated using the z–scaled regulon activity matrix with the selected transcription factors (**Figure 4B**). To determine the overlap between AR, HOXB13 and FOXA1 regulons as per the SCENIC database, a Venn diagram is shown in **Supplementary Figure 10A**.

**Novel cell surface marker detection in AR–positive, AR–negative and NEPC GRNs.** In conjunction with the investigation of known cell surface markers in clinical development (PSMA, STEAP1, STEAP2, CEACAM5, and DLL3), we characterized the most highly expressed cell surface markers in each AR–positive, AR–negative and NEPC regulons. Significantly up–regulated genes for each group (AR–positive, AR–negative and NEPC GRNs) were determined through MAST differential analysis (average $\log_2$FC>0.4, adjusted P–value<0.05). We then restricted to published cell surface markers (**Supplementary Table 16**). A heatmap was generated with the gene expression of the top 15 genes for each group, along with known cell surface markers in clinical development (**Supplementary Figure 10D**).

**DLL3 expression across NEPC and CRPC.** DLL3 expression was analyzed in both CRPC–adenocarcinoma and NEPC cells using both non–imputed and imputed (MAGIC, *k=20, t=1).* Scatter plots were generated for CRPC–adenocarcinoma cells comparing DLL3 expression *versus* CHGB and ASCL1 expression, as well as DLL3 expression *versus* NEPC module score (**Supplementary Figure 10H**). Furthermore, stacked box and density plots of DLL3 non–imputed and imputed expression (MAGIC, *k=20, t=1)* per NEPC regulon were generated (**Supplementary Figure 10E** and **Figure 4E,** respectively)**.** Of note, NEPC module score (used for y–axis in scatter plot) was calculated using the 'AddModuleScore' function (genes defined in **Supplementary Table 13**).

**<u>Intra–Patient Heterogeneity of MSK–HP13 patient</u>**

To address the intra–tumoral heterogeneity of drug target genes, we analyzed each CRPC and NEPC tumor biopsy for variable expression within Phenograph clusters. Specifically, we observed heterogeneity of both PSMA and DLL3 in MSK–HP13. MSK–HP13 tumor cells were subsetted and scaled counts were projected onto 4 PCs. Phenograph clustering (*k*=30) was performed, resulting in 19 clusters. Among 19 clusters which all expressed high AR expression, cluster 11, exhibited high expression of *FOLH1/PSMA,* while the remainder were *FOLH1/PSMA* negative. A heatmap was generated with scaled gene expression of *AR*, *PSMA*, *TACSTD2*, and *DLL3* and the scaled regulon activity of HOXB13 regulon are shown (**Figure 4C** and **Supplementary Figure 10B**). GSEA was performed on $PSMA^{high}$ and $PSMA^{low}$ cells using pre–ranked genes based on MAST differential analysis (refer to '**Identifying DEGs' and 'MAST and GRN pathway annotation')** (**Supplementary Table 12**). Density plots of *AR* and *FOLH1* expression are shown by patient ID in **Supplementary Figure 10C,** along with genes, namely *GATA2*, *HOXB13*, and *SOX4* that display similar expression distributions as *AR* and *FOLH1* in MSK–HP13 upon manual inspection.

Supplementary Figure 1

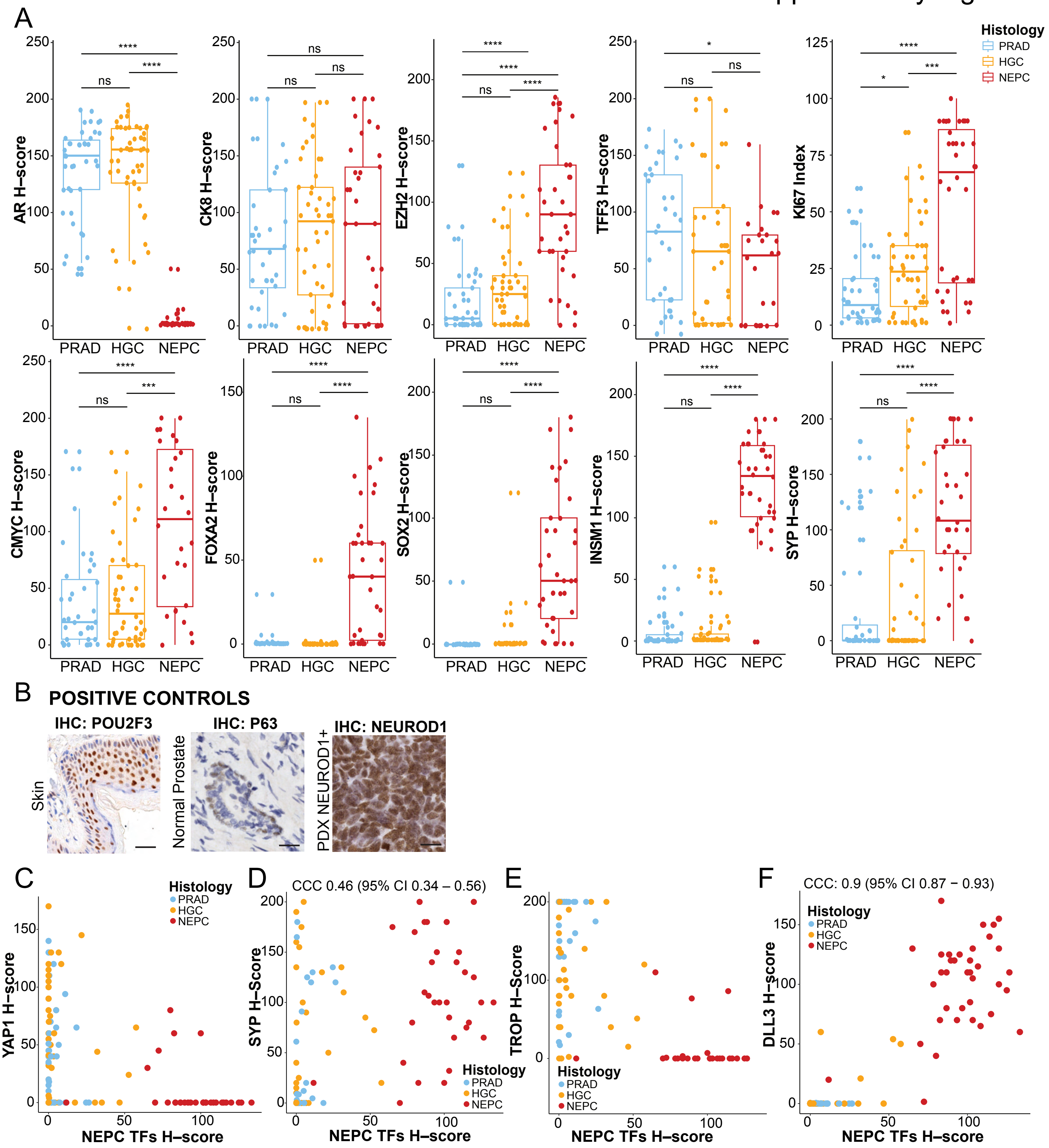

**Supplementary Figure 1. Tissue Microarray of Lineage and Cell Surface Markers in Human CRPC–adenocarcinoma and NEPC. (A)** Boxplot of H–scores of AR**,** CK8, EZH2, TFF3, Ki67 (index score), CMYC, FOXA2, SOX2, INSM1 and SYP and grouped by histology (PRAD, HGC, and NEPC)**.** Significance of H–score distribution between groups was assessed by Wilcoxon signed–rank test. Abbreviations include: not significant (ns), * (<0.05), **(<0.01), ***(<0.001), ****($1x10^{-4}$). **(B)** POU2F3, TP63, and NEUROD1 immunohistochemistry with positive controls in skin, normal prostate (basal cells) and patient derived xenograft—no positivity was detected in TMA cohort. Scale bar: 50µM. **(C–F)** Scatter plot of *H*–scores of YAP1, SYP, TROP2, and DLL3 (y–axis) and NEPC TF H–score (defined as mean H–score of ASCL1, INSM1, and FOXA2) (x–axis), respectively colored by PRAD (light blue), HGC (orange) and NEPC (red). For H–scores with positive linearity, Lin's concordance correlation coefficient (CCC) is noted with 95% confidence intervals.

#### Supplementary Figure 2

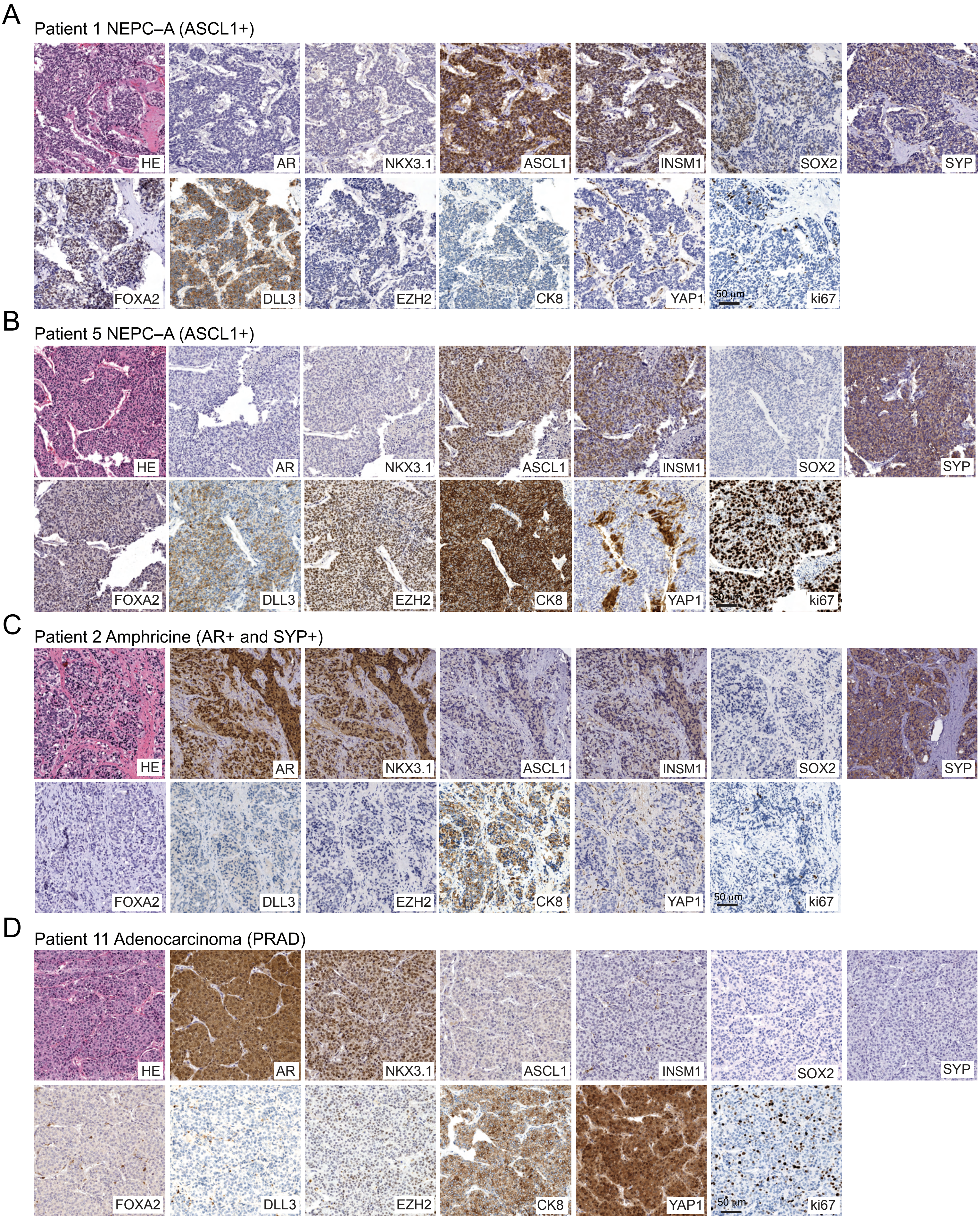

**Supplementary Figure 2. Extended Immunohistochemistry of Tumor Examples from TMA. (A–D)** Hematoxylin and eosin (H&E) and immunohistochemical stains of select tumors from Patient 1 (NEPC–ASCL1/NEPC–A), Patient 5 (NEPC–ASCL1/NEPC–A), Patient 2 (Amphicrine, AR+SYP+ASCL1–), and Patient 11 (Adenocarcinoma, PRAD) are shown. Immunohistochemistry of AR, NKX3.1, ASCL1, INSM1, SOX2, SYP, FOXA2, DLL3, EZH2, CK8, YAP1, and Ki67 are shown; scale bar: 50μM.

Supplementary Figure 3

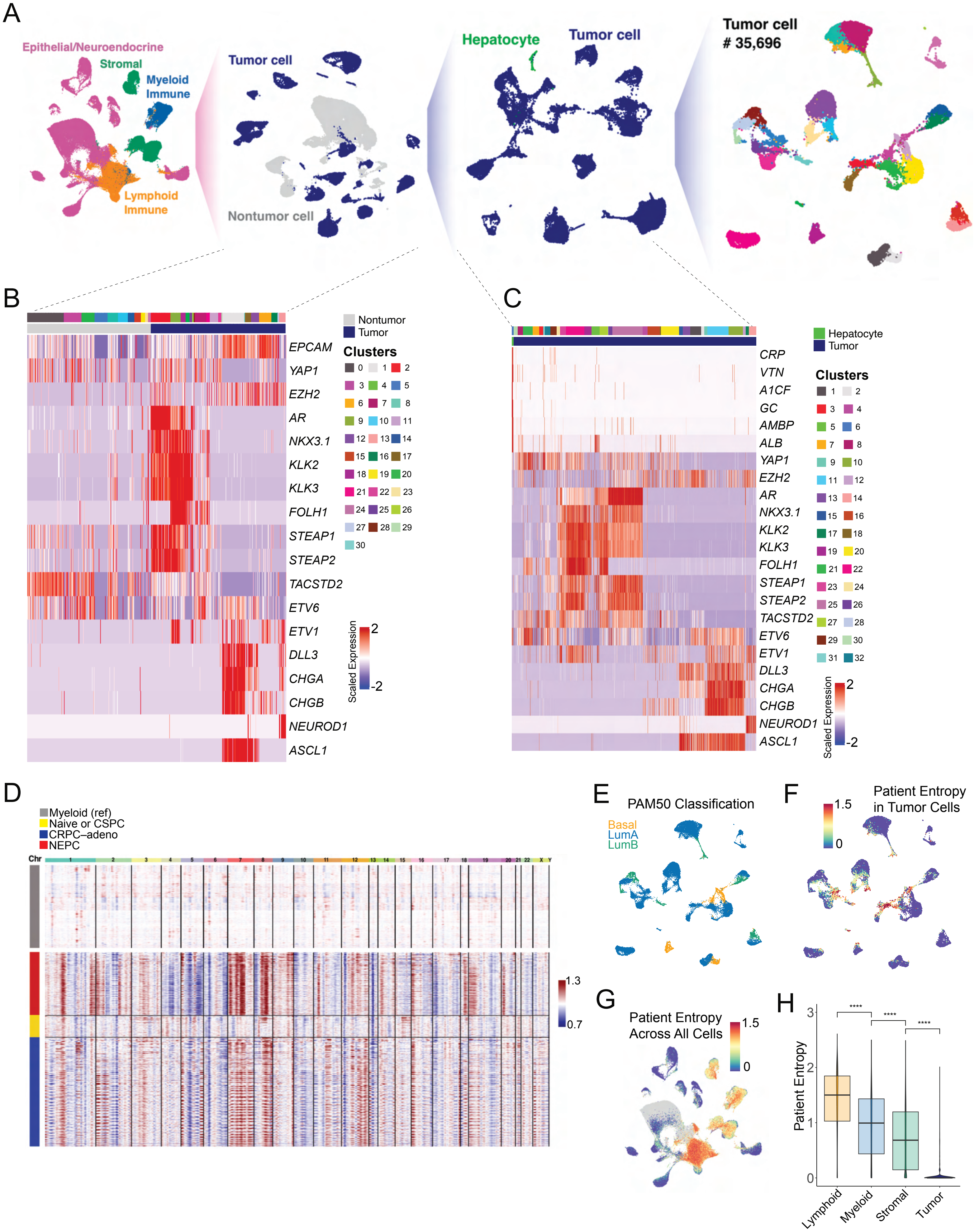

**Supplementary Figure 3. Processing, Entropy, and Tumor Cell Identification in Single Cell RNA–Sequencing. (A)** Schematic for data processing and tumor cell annotation and subsetting (Methods). **(B)** Heatmap of marker gene expression *per cell* (columns) shown for all epithelial/neuroendocrine cells. Cells grouped by clusters (0–30) and as being tumor and nontumor (or benign) as annotated on top of the panel. Z–score of normalized expression (scale: –2 to +2). **(C)** Heatmap of marker gene expression *per cell* (columns) shown for all putative tumor cells. Hepatocyte population was identified as cluster–specific expression of *ALB, CRP, A1CF, AMBP,* which was present in a subset of cells from liver biopsies derived from MSK–HP19, MSK–HP09, MSK–HP17, and MSK–HP18. Cells grouped by clusters (0–32) and as being tumor and hepatocyte as annotated on top of the panel. Z–score of normalized expression (scale: –2 to +2). **(D)** Heatmap of *per* cell copy number variation (CNV) estimated by sliding window approach (scale: modified expression 0.7 to 1.3, Methods) with rows corresponding to cells grouped by cell types (Naïve or CSPC, CRPC–adeno and NEPC with reference set as myeloid cells) and columns corresponding to ordered genomic location. **(E)** Tumor cells scored by PAM50 classification restricted to LumA, LumB, and basal signatures. **(F)** UMAP with inter–patient heterogeneity measured by Shannon entropy of patient frequencies in tumor cells alone. **(G)** To justify batch correction (Methods), entropy–based measure plotted on UMAP of all cells that assesses mixing of normalized expression demonstrating highest entropy in immune compartment, followed by stromal cells, and finally epithelial cells. **(H)** Boxplot of inter–patient heterogeneity measured by Shannon entropy based of patient frequencies. To control for cell sampling, 100 cells were subsampled from each Phenograph cluster (*k*=30) within tumor, immune, and mesenchymal compartments 100 times with replacement (Wilcoxon signed–rank test, Methods). Abbreviations: ****($1x10^{-4}$).

Supplementary Figure 4

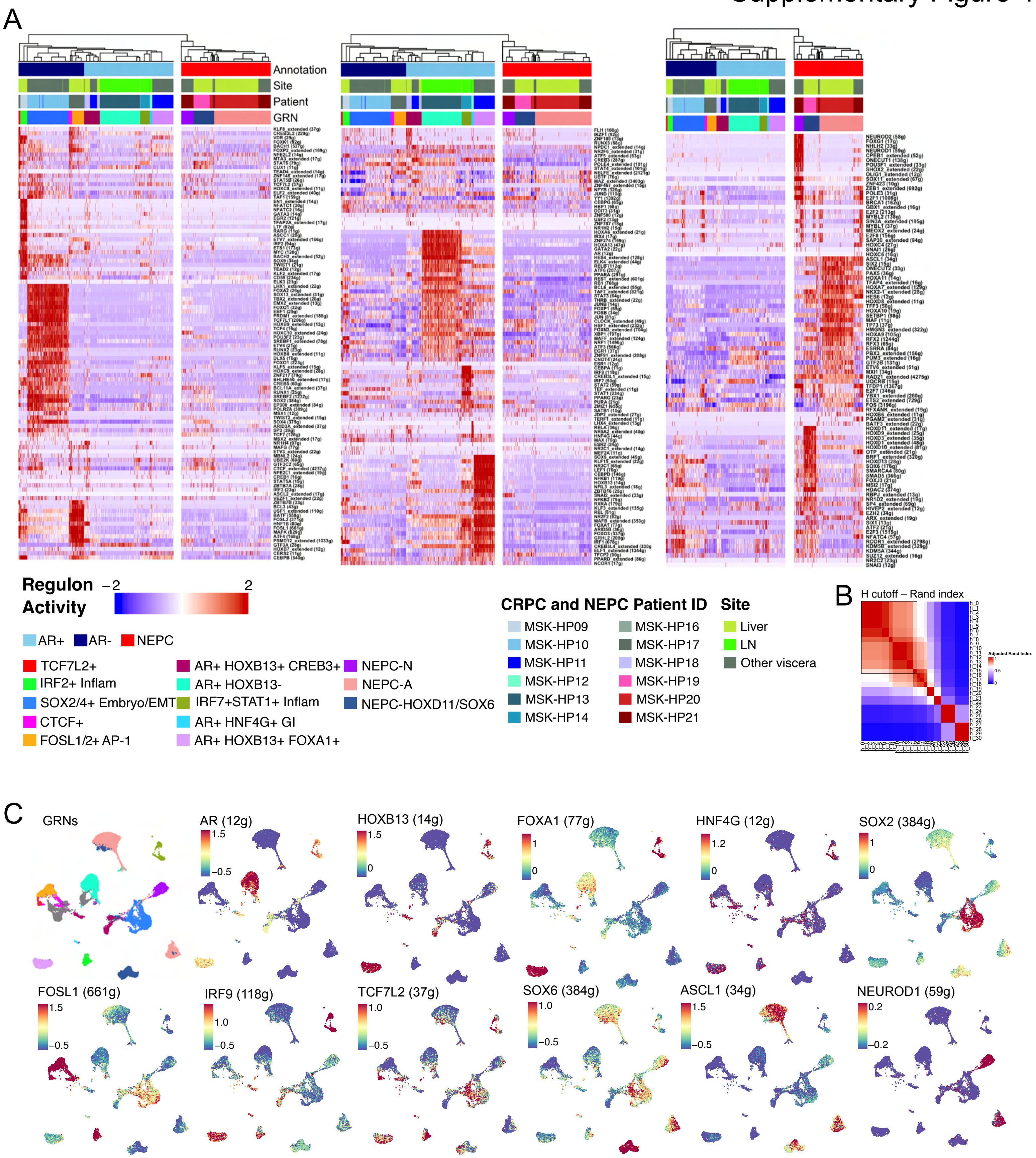

**Supplementary Figure 4. Complete Regulon Analysis in Human CRPC Tumors. (A)** Heatmap of CRPC–adenocarcinoma and NEPC cells (x–axis) and *per* cell scaled regulon activity scores (z–score: –2 to 2) is shown for all TFs (paratheses denotes number of genes within regulon). A dendrogram cutoff of 15 based on adjusted Rand index (shown in (**B**)**,** overall >= 0.65 indicating moderate to high recovery) was used to unbiasedly define the number of gene–regulatory networks (GRNs), yielding 10 and 3 CRPC–adeno and NEPC GRNs, respectively. Regulons were assigned to GRNs based on regulon specificity score (RSS) and ranked for significance using a Student's t–test (Methods). Adenocarcinoma GRNs were labeled based on AR activity (light blue on top panel of heatmap; bracketed by AR–positive GRNs) and without or having low AR activity (dark blue on top panel of heatmap; bracketed by AR–negative GRNs). NEPC regulons are shown (red on top panel of heatmap; bracketed by NEPC GRNs). **(C)** UMAPs are shown colored by GRN (first UMAP on left) or by z–score of regulon activity for select regulons (paratheses denotes number of genes within regulon), which were highlighted in **Figure 2D**.

### Supplementary Figure 5A

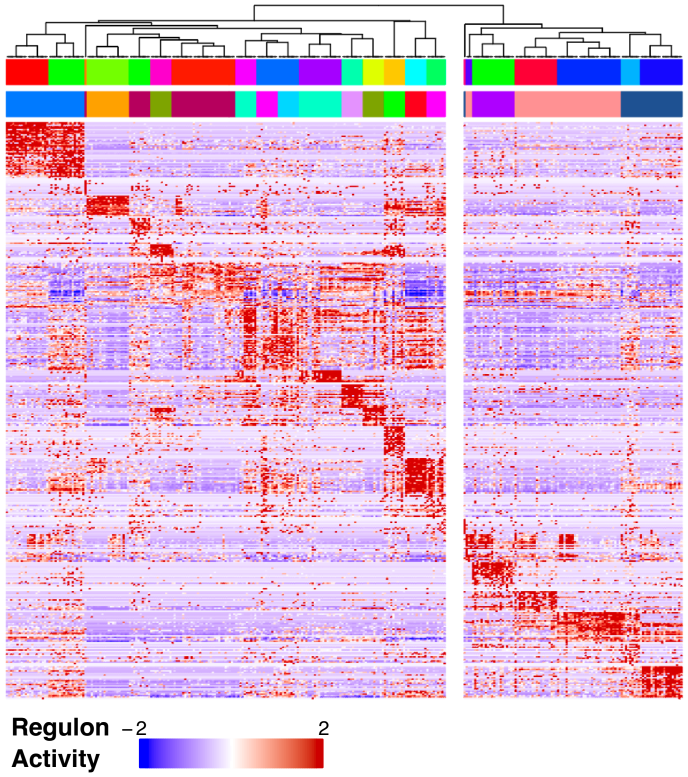


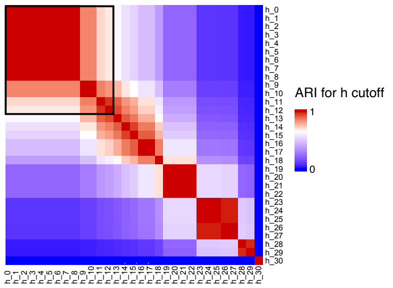


10 random cells for each Phenograph cluser
Random sampling *versus* Human GRNs on complete set
**ARI = 0.69**

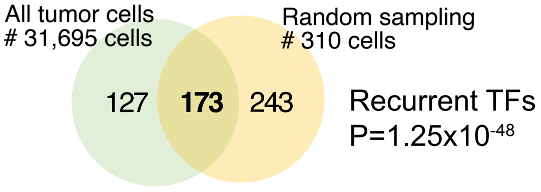


**Random sampling Group** (h =12)

CRPC 1, CRPC 2, CRPC 3, CRPC 4, CRPC 5, CRPC 6, CRPC 7, CRPC 8, CRPC 9, CRPC 10, CRPC 11, CRPC 12, CRPC 13, CRPC 14, CRPC 15, NEPC 1, NEPC 2, NEPC 3, NEPC 4, NEPC 5, NEPC 6, NEPC 7

**Human GRN**

TCF7L2+, CTCF+, IRF2+ Inflammatory, SOX2/4+ Embryo/EMT, FOSL1+ AP-1, AR+ HOXB13+CREB3+, AR+ HOXB13-, AR+ IRF7+ STAT1 Infam, AR+ HNF4G+ GI, AR+ HOXB13+ FOXA1+, NEPC-N, NEPC-A, NEPC-H/SOX6

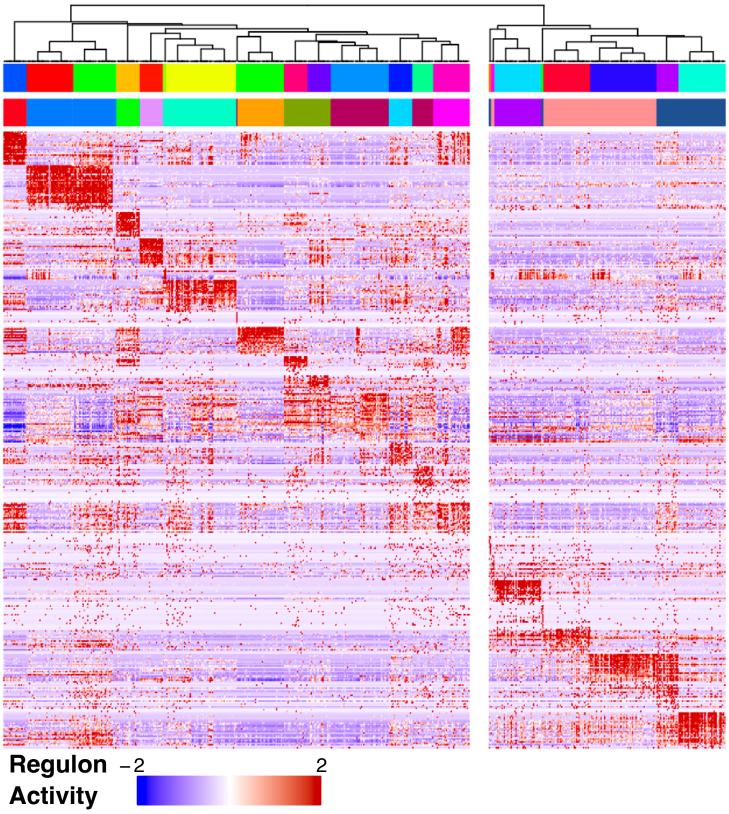


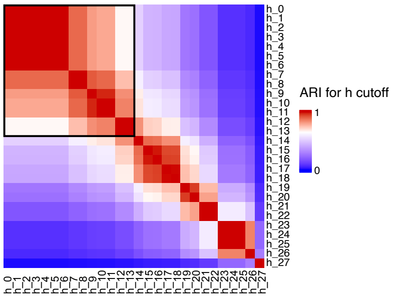


20 random cells for each Phenograph cluser
Random sampling *versus* Human GRNs on complete set
**ARI = 0.75**

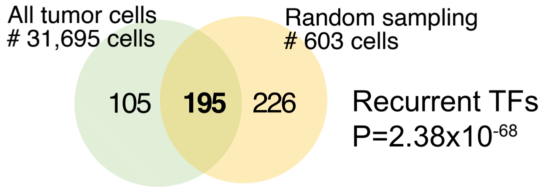


**Random sampling Group** (h =14)

CRPC 1, CRPC 2, CRPC 3, CRPC 4, CRPC 5, CRPC 6, CRPC 7, CRPC 8, CRPC 9, CRPC 10, CRPC 11, CRPC 12, CRPC 13, CRPC 14, CRPC 15, NEPC 1, NEPC 2, NEPC 3, NEPC 4, NEPC 5, NEPC 6, NEPC 7, NEPC 8

**Human GRN**

TCF7L2+, CTCF+, IRF2+ Inflammatory, SOX2/4+ Embryo/EMT, FOSL1+ AP-1, AR+ HOXB13+CREB3+, AR+ HOXB13-, AR+ IRF7+ STAT1 Infam, AR+ HNF4G+ GI, AR+ HOXB13+ FOXA1+, NEPC-N, NEPC-A, NEPC-H/SOX6

#### Supplementary Figure 5B

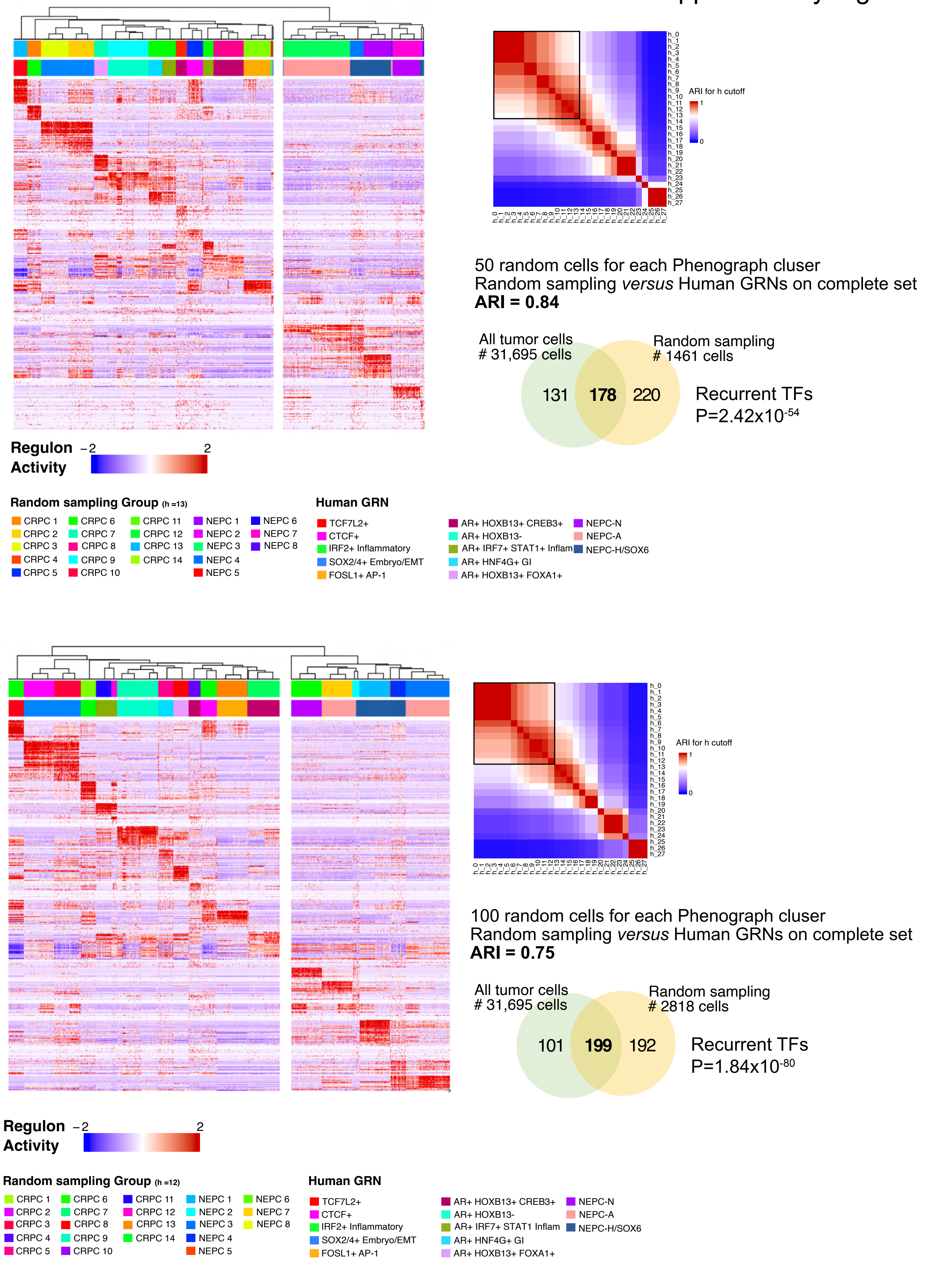

**Supplementary Figure 5A and 5B. Cell Level Robustness Analyses for Human CRPC Regulons.** Robustness analyses were performed by subsampling 10 and 20 (**Supplementary Figure 5A**) or 50 and 100 (**Supplementary Figure 5B**) random cells from each Phenograph cluster (*k=20)*. SCENIC was rerun with analogous methods to the complete dataset (Methods). Shown in heatmap of CRPC–adenocarcinoma and NEPC cells (x–axis) and *per* cell scaled regulon activity scores (z–score: –2 to 2) for all TFs. A dendrogram cutoff of 12 or 13 based on adjusted Rand index to unbiasedly define the number of gene–regulatory networks (GRNs). Regulons were assigned to GRNs based on regulon specificity score (RSS) (Methods). For each run, ARI was calculated between the down–sampled GRN group and the complete GRNs grouping defined by all cells (**bolded**, ARI=0.69–0.84). A Venn diagram also shows the overlapping regulons detected by subsampling compared to our complete run. Significance was assessed by Fisher's exact test.

### Supplementary Figure 6

#### Patient Level Robustness Analysis for GRNs

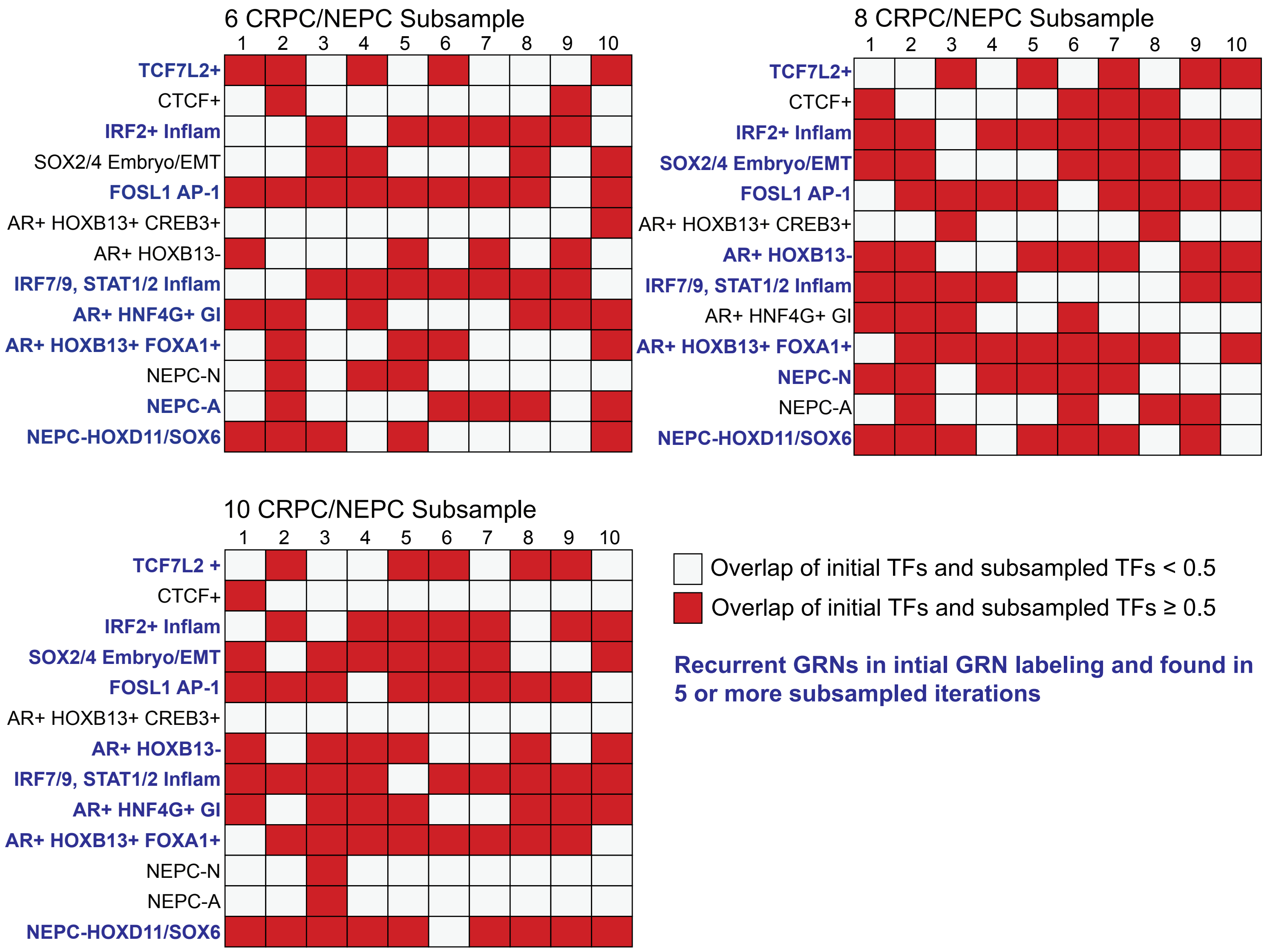

**Supplementary Figure 6. Patient Level Robustness Analyses for Human CRPC Regulons.** Robustness analyses were performed by subsampling 100 cells in 6, 8 and 10 CRPC or NEPC patients (of 13 total CRPC and NEPC patients) and repeated iteratively 10 times with replacement. For each subsampling run, we utilized SCENIC to identify enriched TF/regulons and calculated the proportion of overlapping TFs/regulons between our initial GRNs (Figure 2D) and newly identified subsampled GRNs. For initial and subsampled GRNs that share more than 50% of TFs/regulons, we utilized a Fisher's exact test to determine whether the overlap was significant: P–value $< 0.05$—constituting a shared and recurrent GRN. By subsampling 6, 8, and 10 patients, we identified eight, nine and nine recurrently identified GRNs that significantly overlapped in our initial GRN assignments (and were found in more than 5 of the iterations and bolded and highlighted in blue).

Supplementary Figure 7

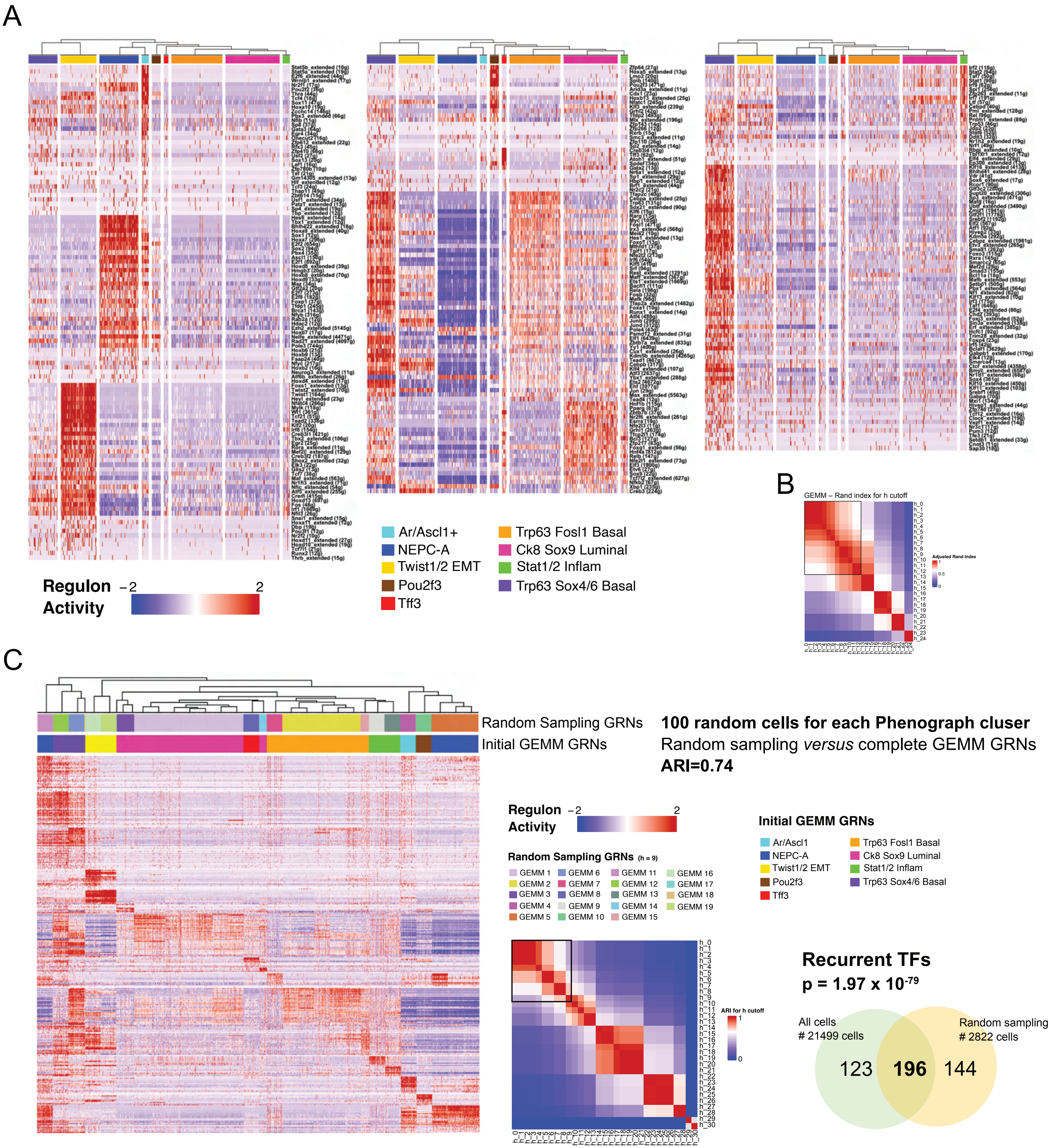

**Supplementary Figure 7. Complete Regulon Analysis in GEMM Tumors and Robustness Analysis (A)** Heatmap of GEMM tumor cells (*N*=21,499) (x–axis) and *per* cell scaled regulon activity scores (z–score: –2 to 2) is shown for all TFs (paratheses denotes number of genes within regulon). A dendrogram cutoff of 12 based on adjusted Rand index (shown in (**B**)) was used to unbiasedly define number of GRNs (groups of regulons), yielding 9 GRNs (groups of regulons) with regulons assigned to GRNs based on regulon specificity score (RSS, Methods). (**C**) Robustness analyses were performed by subsampling 100 random cells from each Phenograph cluster (*k=20)*. SCENIC was rerun with analogous methods to the complete dataset (Methods). Shown in heatmap of GEMM *Gfp*+ cells (x–axis) and *per* cell scaled regulon activity scores (z–score: –2 to 2) for all TFs. A dendrogram cutoff of 9 based on adjusted Rand index to unbiasedly define the number of gene–regulatory networks (GRNs). Regulons were assigned to GRNs based on regulon specificity score (RSS) (Methods). ARI was calculated between the down–sampled GRN group and the complete GRNs grouping defined by all cells (ARI=0.74). A Venn diagram also shows the overlapping regulons detected by subsampling compared to our complete run. Significance was assessed by Fisher's exact test.

### Supplementary Figure 8

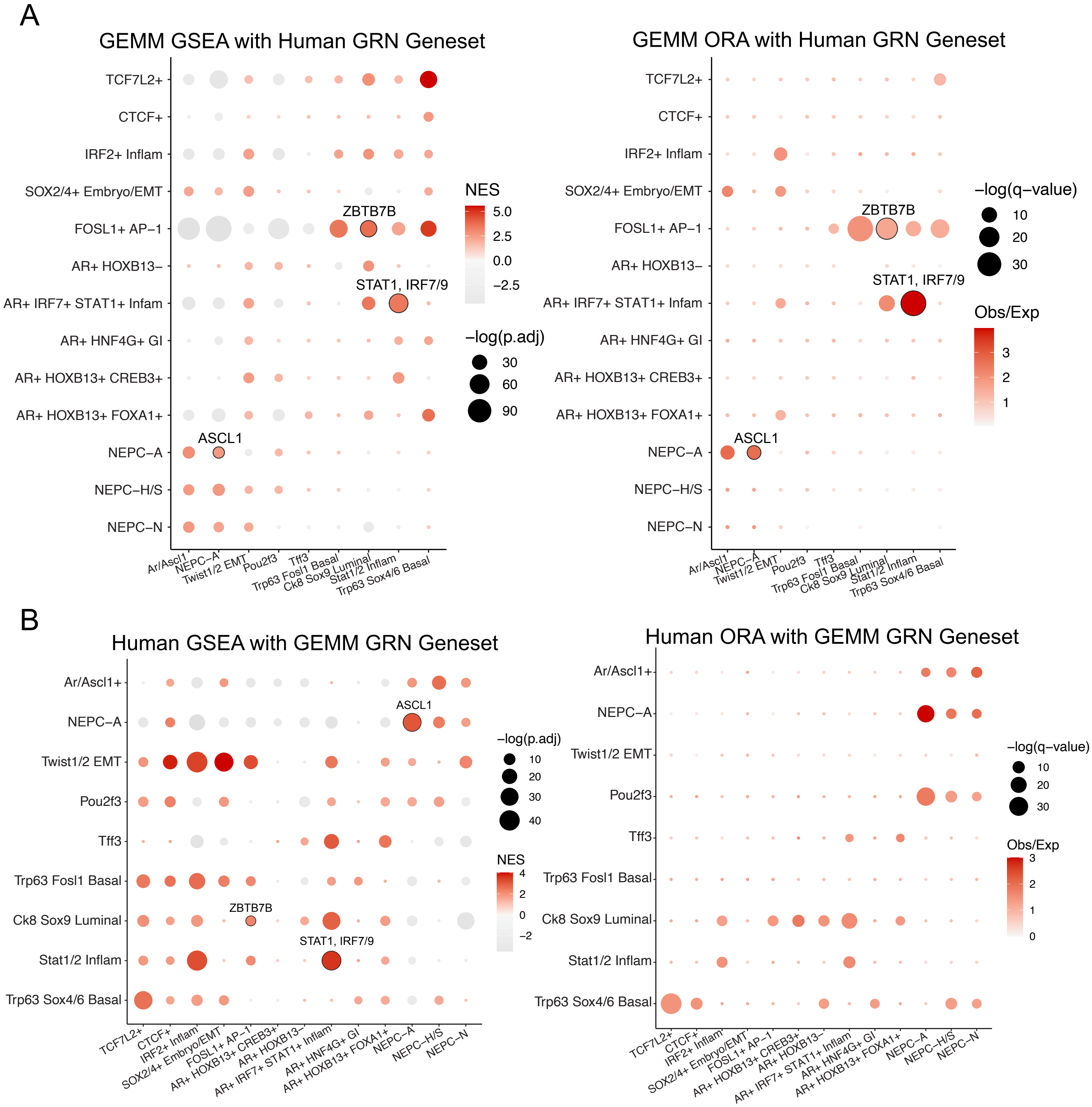

**Supplementary Figure 8. GEMM and Human CRPC/NEPC Overlap.** (**A**) Dot plot for GSEA (left) and over–representation (ORA) (right) analyses for the enrichment of human GRN gene sets in GEMM GRNs. Gene sets were constructed with the top 5 transcription factors (TFs) based on regulon specificity scores (RSS) and their associated target genes for each Human GRN (Methods). GSEA was performed on pre–ranked genes using 10,000 permutations. Gene ranks were calculated based on MAST differential expression (Methods). Dot size denotes the –log(p.adjusted value) and gradient from gray to red is normalized enrichment score (scale: –2.5 to 5). ORA was performed on DEGs with average $\log_2$FC > 0.25, adjusted p–value < 0.05. Dot size denotes the –log(q–value) and gradient from gray to red is observed over expected ratio (scale: 0 to 3). Shared GRN pairs having significant enrichment in all iterative tests are outlined in black with the shared top TF labeled (Methods). (**B**) Dot plot for GSEA (left) and over–representation (ORA) (right) analyses for the enrichment of GEMM GRN gene sets in Human GRNs. Gene sets were constructed with the top 5 transcription factors (TFs) based on regulon specificity scores (RSS) and their associated target genes for each GEMM GRN (Methods). GSEA was performed on pre–ranked genes using 10,000 permutations. Gene ranks were calculated based on MAST differential expression (Methods). Dot size denotes the –log(p.adjusted value) and gradient from gray to red is normalized enrichment score (scale: –2 to 4). ORA was performed on DEGs with average $\log_2$FC > 0.25, adjusted p–value < 0.05. Dot size denotes the –log(q–value) and gradient from gray to red is observed over expected ratio (scale: 0 to 3). Shared GRN pairs having significant enrichment in all iterative tests are outlined in black with the shared top TF labeled (Methods).

### Supplementary Figure 9

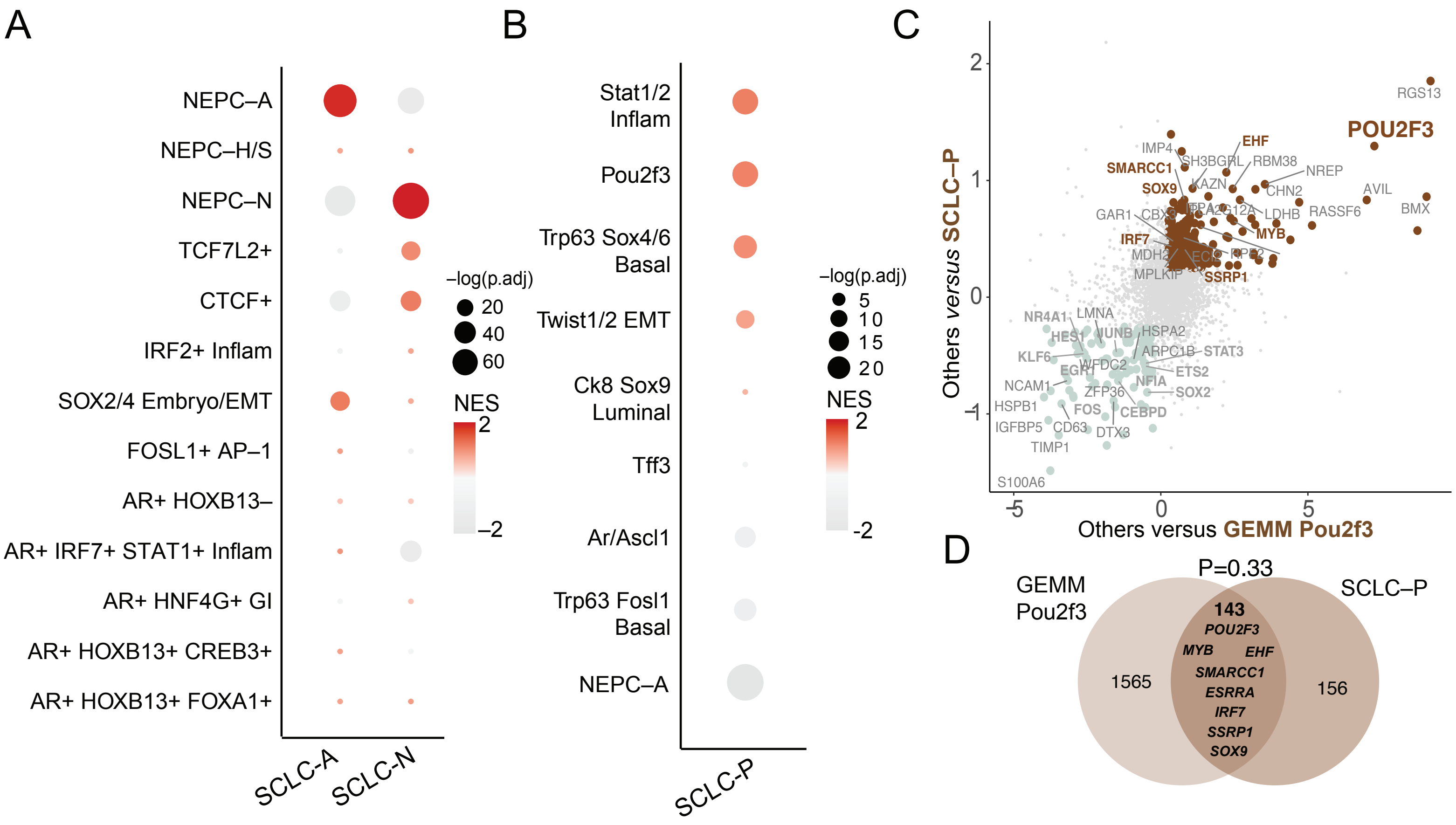

**Supplementary Figure 9.** ***Ascl1, Neurod1, and Pou2f3*** **Overlap in SCLC and NEPC (A)** Dot plot for GSEA analysis for the enrichment of human GRN gene sets in SCLC subtypes (SCLC–A and SCLC–N). Gene sets were constructed with the top 5 transcription factors (TFs) based on regulon specificity scores (RSS) and their associated target genes for each Human GRN (Methods). GSEA was performed on pre–ranked genes using 10,000 permutations. Gene ranks were calculated based on MAST differential expression (Methods). Dot size denotes the –log(p.adjusted value) and gradient from gray to red is normalized enrichment score (scale: –2 to 2). **(B)** Dot plot for GSEA analysis for the enrichment of GEMM GRN gene sets in SCLC–P subtype. Gene sets were constructed with the top 5 transcription factors (TFs) based on regulon specificity scores (RSS) and their associated target genes for each GEMM GRN (Methods). GSEA was performed on pre–ranked genes using 10,000 permutations. Gene ranks were calculated based on MAST differential expression (Methods). Dot size denotes the –log(p.adjusted value) and gradient from gray to red is normalized enrichment score (scale: –2 to 2). (**C**) SCLC–P *vs.* rest (shown on y–axis) or GEMM *Pou2f3 vs.* rest (shown on x–axis) were compared using MAST and the $\log_2$FC for each gene is shown on the scatter plot. Genes with $\log_2$FC > 0.4 (and $p_{adj}<0.05$) are labeled with TFs noted in brown for being enriched in both SCLC–P and GEMM Pou2f3, respectively. (**D**) Venn diagram shows the overlap of top DEGs (average $\log_2$FC > 0.4, adjusted p–value < 0.05) shared between SCLC–P and GEMM *Pou2f3*. A Fisher's exact test was used for significance of overlap.

### Supplementary Figure 10

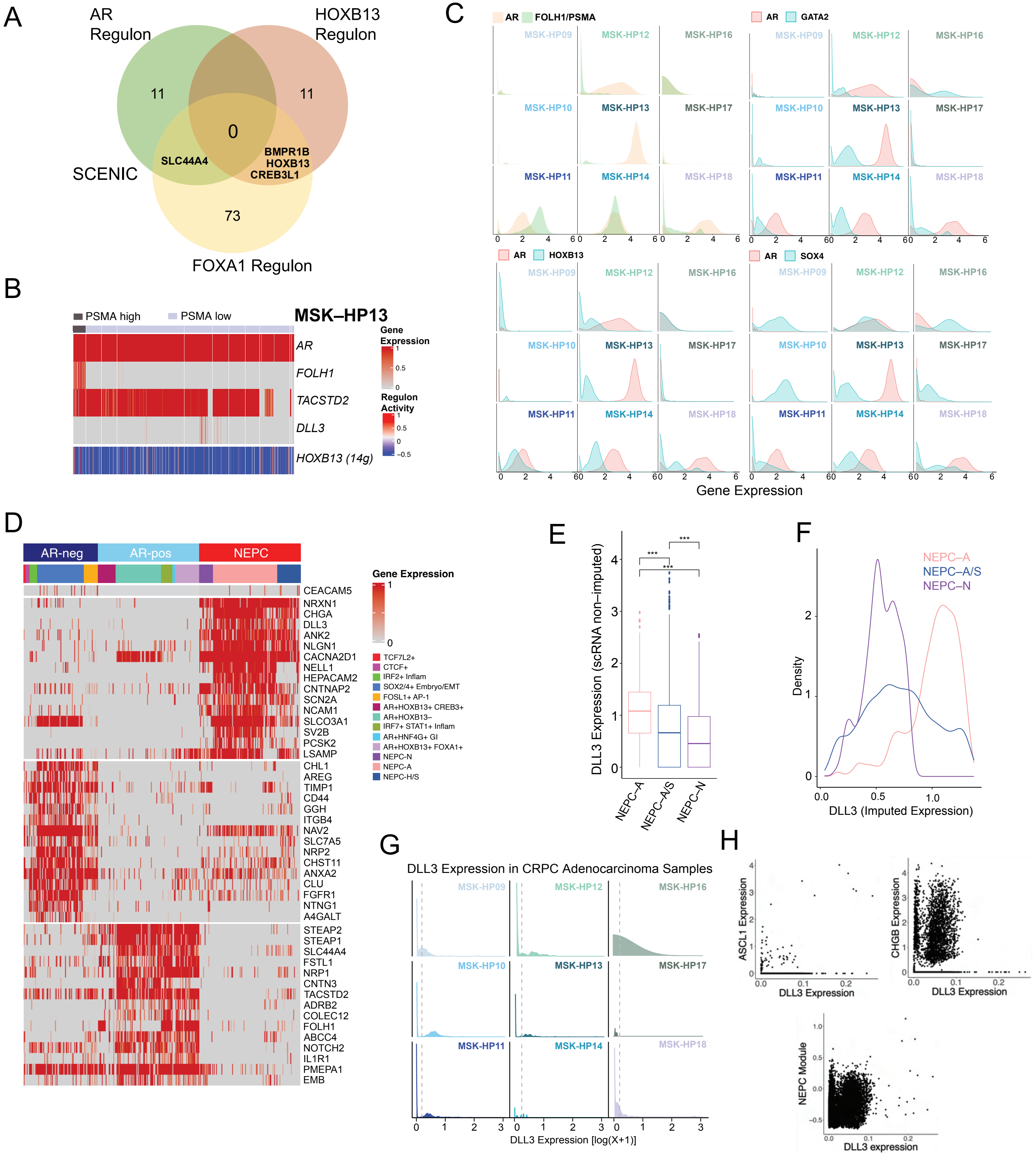

**Supplementary Figure 10. *FOLH1*, *DLL3*, and Novel Cell Surface Marker Detection in GRNs.** **(A)** Overlap of AR, HOXB13, FOXA1 regulon (and target genes) per SCENIC data (cis target database). **(B)** Heatmap of scaled expression (z–score) is shown for *AR*, *FOLH1*, and *TACSTD2* genes, and *HOXB13* regulon activity in MSK–HP13. **(C)** Density plots *per* CRPC–adenocarcinoma sample of *AR* and *FOLH1/PSMA* expression (note MSK–HP13 with discordant *AR* and *FOLH1/PSMA* expression), along with genes that follow *FOLH1/PSMA* distribution as in MSK–HP13, namely *GATA2*, *HOXB13*, and *SOX4*. (**D**) Heatmap of normalized gene expression (0 to 1) showing existing and novel cell surface markers in AR–positive, AR–negative and NEPC GRNs. **(E)** Non–imputed DLL3 expression log(X+1) grouped by NEPC regulons. **(F)** Density plot of DLL3 imputed expression (MAGIC, *k=20, t=1*) by NEPC regulon. **(G)** Density plots *per* CRPC–adenocarcinoma sample of non–imputed DLL3 expression log(X+1) suggesting positivity in a subset of cells. **(H)** Scatter plots of non–imputed DLL3 expression log(X+1) of CRPC–adenocarcinoma cells with non–imputed expression [log(X+1) of *CHGB* or *ASCL1,* or NEPC module score (Methods).
